# *Plasmodium falciparum* Atg18 localizes to the food vacuole via interaction with the multi-drug resistance protein 1 and phosphatidylinositol 3-phosphate

**DOI:** 10.1101/2020.10.02.323600

**Authors:** Renu Sudhakar, Divya Das, Subramanian Thanumalayan, Somesh Gorde, Puran Singh Sijwali

**Author notes:** **Corresponding author:** Puran Singh Sijwali. **Abbreviations.** *Plasmodium falciparum* Atg18: PfAtg18; WIPI: WD-repeat protein interacting with phosphoinositides; PROPPIN: β-propellers that bind polyphosphoinositides; PI3P: Phosphatidylinositol 3-phosphate; PI3K: Phosphoinositide 3-kinase; Vps: Vacuolar Protein Sorting; LC3: Microtubule-associated protein 1A/1B-light chain 3; LIR: LC3-interacting region; AIM: Atg8-family interacting motif; MDR-1: multidrug resistance protein 1; phagophore assembly site: PAS; UTR: Untranslated region; GST: Glutathione S-transferase; DHFR: Dihydrofolate Reductase; CDD: C-terminal Destabilizing Domain; IC_50_: half maximal inhibitory concentration; PDB: Protein Data Bank; PCV: packed cell volume; BSD: blasticidin.

## Abstract

Autophagy is a lysosome-dependent degradative process involving over 35 Atg proteins. The autophagy repertoire in malaria parasites is limited and does not appear to be a major degradative process. To better understand the autophagy process, we investigated *Plasmodium falciparum* Atg18 (PfAtg18), a PROPPIN family protein, whose members like *S. cerevisiae* Atg18 (ScAtg18) and human WIPI2 are essential for autophagy. Wild type and mutant PfAtg18 were expressed in *P. falciparum* and assessed for localization, the effect of various inhibitors and antimalarials on PfAtg18 localization, and identification of PfAtg18-interacting proteins. PfAtg18 is expressed in asexual erythrocytic stages and localized to the food vacuole, which was also observed with other *Plasmodium* Atg18 proteins, indicating that food vacuole localization is a conserved feature. Interaction of PfAtg18 with the food vacuole-associated PI3P is essential for localization, as PfAtg18 mutants of PI3P-binding motifs neither bound PI3P nor localized to the food vacuole. Interestingly, ScAtg18 showed complete cytoplasmic localization despite binding with PI3P, indicating additional requirement for PfAtg18 localization. The food vacuole multi-drug resistance protein 1 (MDR1) was consistently identified in the PfAtg18 immunoprecipitate, and also interacted with PfAtg18. In contrast to PfAtg18, ScAtg18 did not interact with the MDR1, which, in addition to PI3P, could play a critical role in localization of PfAtg18. Chloroquine and amodiaquine greatly affected PfAtg18 localization, suggesting that these quinolines target PfAtg18 or the proteins that might be involved in its localization. Thus, PI3P and MDR1are critical mediators of PfAtg18 localization, and PfAtg18 may modulate MDR1 activity.

## Introduction

Malaria parasites undergo multi-stage development in diverse intracellular and extracellular environments. Several of these stages are specialized for invasion, degradation of host cellular contents to obtain nutrients, generation of trafficking system for import and export, and extensive reorganization of intracellular organelles to meet stage-specific needs. Since autophagy performs both degradative and biosynthetic functions, and several autophagy proteins have key roles in endosomal transport and organelle reorganization, investigation of autophagy during parasite development is warranted.

Multiple independent studies focussing on Atg8 proteins of human malaria parasite *P. falciparum* and mouse malaria parasite *P. berghei* indicate a limited autophagy repertoire in *Plasmodium* species [1–6]. It is not clear whether autophagy performs a degradative function in malaria parasites, as the majority of Atg8 puncta remain outside the food vacuole, a lysosome-like organelle wherein bulk degradation of haemoglobin occurs [5]. Autophagy may have a role in endosomal transport to the food vacuole, as some Atg8 puncta were observed near or within the food vacuole [3]. A role of autophagy in the biogenesis and/or the processes associated with apicoplast, a nonphotosynthetic plastid remnant that is essential for biosynthesis of isoprenoid precursors in *Plasmodium*, has been supported by colocalization of Atg8 with apicoplast [1–6], adverse effect on microneme clearance and apicoplast proliferation upon overexpression of Atg8 during *P. berghei* liver stage development [7], and loss of apicoplast in Atg8 knock-down parasites [8, 9]. Although mechanistic details of the association between Atg8/autophagy pathway and apicoplast remain to be uncovered, autophagy may mediate transport of biomolecules and membranes to apicoplast. Knock-down of Atg7 and chemical inhibition of Atg3, two essential enzymes of the Atg8 conjugation system, also impaired parasite development, indicating that Atg8 conjugation system is indispensable [10, 11]. Autophagy pathway has also been associated with resistance to chloroquine, as treatment with the LD_50_ concentration of chloroquine significantly altered the distribution of Atg8 puncta in chloroquine sensitive strain compared to the resistant strain of *P. falciparum* [12]. Thus, multiple lines of data conclude to an atypical, but essential, autophagy pathway in malaria parasites, which could offer attractive targets for new chemotherapeutic interventions.

All the reports in uniformity indicate the presence of Atg8 puncta in malaria parasites, which resemble autophagosomes, the hallmark of autophagy. The formation of autophagosome requires regulated and orderly action of multiple proteins, including the Vps34 complex, Atg18/WIPI4-Atg2 complex, Atg12-Atg5-Atg16 complex and its conjugation system, and the Atg8 conjugation system [13–15]. Upon induction of autophagy, the phosphatidylinositol 3-kinase complex (Vps34/Atg6/Vps15/Atg14L) produces phosphatidylinositol 3-phosphates (PI3P) at the endoplasmic reticulum in mammalian cells and at a peri-vacuolar site in yeast, leading to the generation of PI3P-rich structures, called omegasomes or cradles [16, 17]. The omegasome gives rise to phagophore that is elongated and expanded into autophagosome; this process requires Atg9 for adding membranes to the growing structure, and Atg8/LC3 and Atg12 conjugation systems.

Specific PROPPIN family proteins that bind PI3P are recruited at the omegasomes/cradles. Atg18, Atg21 and Hsv2 of yeast and WIPI1-4 of humans are PROPPIN-family proteins, which contain WD40 repeats and a conserved “FRRG” motif that mediates binding to PI3P and phosphatidylinositol 3,5-bisphosphate [18–23]. *S. cerevisiae* Atg18 (ScAtg18) is essential for macroautophagy and cytoplasm-to-vacuole targeting (Cvt) pathway [24, 25], Atg21 is required for Cvt pathway but not for starvation-induced autophagy [26–28], and Hsv2 appears to have a role in micronucleophagy [26, 29, 30]. The Atg18 proteins of *S. cerevisiae* and *P. pastoris* are also required for pexophagy [31]. Atg18 associates with Atg2, forming the Atg18-Atg2 complex, which is recruited to the phagophore assembly site (PAS) through interaction of Atg18 with PI3P [32, 33]. Atg18-Atg2 association is essential for autophagosome biogenesis [34], and is also required for recycling of Atg9 [35]. WIPI1 and WIPI2 have also been shown to bind PI3P-rich sites on the ER, phagophore and autophagosome through interaction with PI3P [36]. WIPI2 has been shown to recruit the Atg12-Atg5-Atg16L complex at the phagophore, which facilitates conjugation of LC3 to phosphatidylethanolamine on the phagophore membrane [36, 37]. In *S. cerevisiae*, Atg21 contributes to macroautophagy via recruitment of Atg12-Atg5-Atg16L complex and Atg8 at the PAS [38, 39]. A complex of ScAtg18 on the vacuole membrane regulates PI3P and PI3,5P2 levels, which has been shown to be important for maintaining vacuole morphology [40–42]. ScAtg18 and ScAtg21 also localize to endosomes, but the functional significance of this association is not known [29]. Several autophagy proteins that are involved in induction (Atg1), initiation (Vps34), and assembly (Vps34, Atg8, Atg9) of PAS have been shown to colocalize with Atg18 at a peri-vacuolar membrane compartment during starvation condition in yeast, suggesting that this compartment could be a site of vesicle formation or membrane source [43].

Both PI3P and PI3K/Vps34 have been reported in *P. falciparum* erythrocytic stages [44, 45]. However, malaria parasites appear to lack several autophagy proteins that are essential for autophagosome formation, including the subunits of Vps34 complex (Atg6 and Atg14), Atg2, Atg10 and Atg16. Interestingly, a recent report showed that *Plasmodium* Atg12 and Atg5 form a noncovalent complex, which could mediate lipidation of Atg8 [46]. Nonetheless, how and where are Atg8 puncta generated in malaria parasites remains unclear.

All *Plasmodium* species encode for a highly conserved Atg18 homolog [5]. An earlier study showed that *P. falciparum* Atg18 (PfAtg18) is expressed in asexual erythrocytic stages and localizes to vesicular structures in the cytoplasm, which also colocalized with the food vacuole and apicoplast [47]. This study also showed that PI3P is critical for localization of PfAtg18 to vesicular structures, and knock-down of PfAtg18 caused reduced conjugation of Atg8 to membranes and the loss of apicoplast, thereby indicated a crucial role of PfAtg18 in regulation of apicoplast and autophagy. Using colocalization and inhibition approaches, a recent study reported that PfAtg18 has a role in the food vacuole dynamics and autophagy pathway [48]. A mutation in the PfAtg18 gene has been implicated in artemisinin resistance and immunoprecipitate of *P. falciparum* using an artemisinin probe pulled-down Atg18 [49, 50], suggesting an association with artemisinin action. A single nucleotide polymorphism leading to Thr38Ile change in PfAtg18 has recently been shown to be associated with resistance to artemisinin derivatives and better survival of mutant parasites in nutrient-deprivation conditions than wild type parasites [51]. However, a link between Atg18/autophagy pathway and the associated drug resistance is not clear.

We hypothesized that investigation of PfAtg18 could provide insights into the formation of Atg8 puncta and the autophagy-associated drug resistance. Our data indicate that food vacuole localization is a conserved feature of *Plasmodium* Atg18, which is mediated via interaction with PI3P and the food vacuole membrane multi-drug resistance protein 1 (MDR1). The food vacuole localization of PfAtg18 was altered upon treatment of parasites with chloroquine and amodiaquine.

## Experimental procedures

### Materials

All the biochemicals were from Sigma or Serva unless otherwise mentioned. The parasite culture reagents were from Lonza and Thermo Fisher Scientific. Restriction enzymes and DNA modifying enzymes were from New England Biolabs and Thermo Fisher Scientific. DNA isolation kits were from QIAGEN and MACHEREY-NAGEL. Dialysis membrane, ProLong Diamond Antifade Mountant, Hoechst, DAPI, LysoTracker® Red DND-99 and SuperSignal Chemiluminescent substrates were from Thermo Fisher Scientific. Antibodies were from Sigma and Thermo Fisher Scientific. WR99210 was a kind gift from David Jacobus (Jacobus Pharmaceutical, Princeton, USA). *P. falciparum* 3D7 and D10 strains, *P. berghei* ANKA strain and *P. knowlesi* (strain H) genomic DNA were obtained from the Malaria Research and Reagent Reference Resource centre (MR4). *P. vivax* genomic DNA was a kind gift from Dr. Kailash C Pandey of the National Institute of Malaria Research, New Delhi, India. Human blood was collected after obtaining written informed consent from all the subjects, processed according to the protocols (IEC-38/2015, IEC-38-R1/2015, IEC-38-R2/2015 and IEC-38-R3/2015) approved by the Institutional Ethics Committee of Centre for Cellular and Molecular Biology, and studies abide by the Declaration of Helsinki principles. Animals were housed in cabin type isolators at standard environmental conditions (22-25°C, 40-70% humidity, and 12:12 hour dark/light photoperiod). All animal experiments were carried out according to the protocols (13/2014, 5/2015, 38/2016, 4/2017 and 28/2018) approved by the Institutional Animal Ethics Committees (IAEC) of Centre for Cellular and Molecular Biology.

### Sequence analysis and homology modelling

Sequences of selected characterized Atg18/WIPI proteins were obtained from the UniProt database (PaAtg18: Q5QA94; KmHsv2: J3QW34; DmAtg18a: Q9VSF0; ScAtg18: P43601; CeAtg18: O16466; DrWIPI2: F1QLJ9; HsWIPI2: Q9Y4P8), and aligned with PfAtg18 (PF3D7_1012900) using T-Coffee (http://tcoffee.crg.cat/apps/tcoffee/do:mcoffee). The PfAtg18 sequence was analysed for conserved motifs and domains using the iLIR and WD40 repeat protein Structure Predictor (WDSP) programs [52, 53]. PDB was searched to identify the closest structural homologs of PfAtg18, which were used as templates (*Kluyveromyces marxianus* Hsv2, PDB ID: 3VU4; *Pichia angusta* Atg18, PDB ID: 5LTD) for generating the modelled structure of PfAtg18 using the SWISS-MODEL server [54]. The model was viewed and edited using the PyMOL Molecular Graphics System (version 1.7.6.0 Schrödinger, LLC). Transmembrane helices in PfMDR1 (PF3D7_0523000.1) were predicted using the TMHMM 2.0 server (http://www.cbs.dtu.dk/services/TMHMM-2.0/).

### Parasite culture, isolation of nucleic acids and cDNA synthesis

*P. falciparum* 3D7 and D10 strains were grown in RPMI 1640-albumax medium with human erythrocytes at 2% haematocrit (RPMI 1640 supplemented with 2 g/l sodium bicarbonate, 2 g/l glucose, 25 µg/ml gentamicin, 300 mg/l glutamine, 100 µM hypoxanthine, 0.5% albumax II) [55]. Synchronization of parasites was achieved by treatment with sorbitol when the majority of parasites were in the ring stage, and parasites were isolated from infected RBCs by lysis with saponin [5, 56]. Parasite pellets were used immediately or stored at −80°C till further use. Genomic DNA was isolated from late trophozoite/schizont stage parasites using the Puregene Blood Core kit according to the manufacturer’s protocol. For RNA isolation, the parasite pellet (∼50 µl PCV) was resuspended in 300 µl Trizol solution, mixed with 200 µl chloroform, centrifuged to separate the upper layer, which was mixed with 300 µl isopropanol to precipitate total RNA. The RNA pellet was washed with 300 µl of 70% ethanol, resuspended in RNAse-free water, and stored at −80°C until further use. cDNA was made from the total RNA using the SuperScript® III First-Strand Synthesis System as instructed by the manufacturer. Briefly, 5 µg of gDNA-free total RNA was mixed with random hexamer primers and dNTPs, heated at 65°C for 5 min, followed by immediate cooling in ice. 10 µl of this mixture was incubated with Reverse Transcriptase and RT buffer at 30°C for 10 min, followed by at 50°C for 1 hour. The reaction was terminated by incubating the mixture at 85°C for 5 min, and 1-2 µl of this sample was used in PCR.

For *P. berghei* parasites, 3-6 weeks old BALB/c mice were infected with a frozen stock of *P. berghei* ANKA and infection was monitored regularly by observing Giemsa stained blood smears of the tail snips of infected mice. The infected mice were euthanized at 10-15% parasitemia, blood was collected in Alsever’s solution (2.05% glucose, 0.8% sodium citrate, 0.055% citric acid and 0.42% sodium chloride) by cardiac puncture, parasites were purified by saponin lysis method, and the parasite pellets were processed for isolation of genomic DNA using the Puregene Blood Core kit according to the manufacturer’s protocol.

### Expression and purification of recombinant proteins

The PfAtg18 C-terminal coding region (PfAtg18ct: 571-1143 bps) was amplified from *P. falciparum* cDNA using PfATG18-F2/Atg18expR primers (Table S2), and cloned into the pCR2.1 vector to obtain pCR2.1-PfAtg18ct. The insert was subcloned into pET32a at BamHI/HindIII sites to obtain pET32a-PfAtg18ct, which was transformed into BL21-CodonPlus(DE3)-RIL cells. pET32a-PfAtg18ct would express Thioredoxin/His-PfAtg18ct (Trx/His-PfAtg18ct) fusion protein, which was purified from IPTG induced cell under denaturing conditions. Briefly, the induced cell pellet was lysed in urea buffer (8M urea, 50 mM NaH_2_PO_4_, 500 mM NaCl, pH 8.0; 5 ml buffer/g pellet), sonicated using the SONICS Vibra Cell Ultrasonic Processor (9 secs pulses at 20% amplitude for 4 min), centrifuged, and the supernatant was separated. Imidazole (10 mM final), Triton-X 100 (0.5% final) and β-ME (5 mM final) were added to the supernatant, and incubated with Ni-NTA agarose resin (0.25 ml slurry/g weight of the initial pellet for 30 min at room temperature). The resin was washed with 50x column volume of wash buffer 1 (urea buffer with 30 mM imidazole, 0.5% Triton-X 100 and 5 mM β-ME) and wash buffer 2 (urea buffer with 50 mM imidazole). The bound proteins were eluted with elution buffer (250 mM imidazole in urea buffer) and assessed for purity by SDS-PAGE. Elution fractions containing pure protein were pooled and refolded by dialyzing in a 10 kDa cut off dialysis tubing against the refolding buffer (20 mM Tris pH 7.5, 1 mM EDTA, 0.5 mM DTT, 50 mM NaCl, 10% glycerol) at 4^°^C for 18 hrs with one change of buffer after 14 hours. The dialysed protein was concentrated in a 10 kDa cut off Amicon Ultra-15 (Millipore), quantitated using BCA, and used for immunizing rats.

A synthetic and codon optimised PfAtg18 gene was purchased from GenScript as a pUC57-PfAtg18syn plasmid, which contained PfAtg18 coding sequence flanked by the NotI/EcoRI sites. The PfAtg18syn insert was subcloned into the pGEX-6P-1 at EcoRI/NotI sites to generate pGEX/PfAtg18syn. PfAtg18 mutants of PI3P-binding motifs (PIPm, FRRG mutated to FAAG; ALCAm, WLCL mutated to ALCA) were generated by recombination PCR. PIPm was amplified from the pUC57-PfAtg18syn plasmid as two fragments using FAAG Syn F/M13R and M13F/FAAG Syn R primers, and the two fragments were recombined using M13F/M13R primers. Similarly, ALCAm was amplified from the pUC57-PfAtg18syn plasmid using the primer sets ALCA Syn F/M13R and M13F/ALCA Syn R, and the two fragments were recombined. The PIPm and ALCAm fragments were digested with EcoRI/NotI sites and cloned into similarly digested pGEX-6P-1 vector to obtain pGEX/PIPm and pGEX/ALCAm, respectively. The ScAtg18 coding region contains internal BamHI and BglII sites, which were sequentially eliminated without affecting the encoded-amino acids by recombination PCR. For elimination of BamHI site, ScAtg18 was amplified from the genomic DNA of wild type BY4741 strain as two fragments using SCATG18EXPF/SCATG18MBR and SCATG18MBF/SCATG18EXPR primer sets. The two fragments were recombined using the primers SCATG18EXPF/SCATG18EXPR and cloned into the pGT-GFPbsc vector at BamHI/XhoI sites to obtain pGT-GFPScAtg18-Bm. The pGT-GFPbsc plasmid has been described previously [57]. For elimination of BglII site, pGT-GFPScAtg18-Bm was used as a template using primer sets SCATG18EXPF/SCATG18MBGR and SCATG18MBGF/SCATG18EXPR. The two fragments were recombined using SCATG18EXPF/SCATG18EXPR primers and cloned into the pGT-GFPbsc plasmid at BamHI/XhoI sites to obtain pGT-ScAtg18-BmBgm. The ScAtg18BmBgm insert was subcloned into the pGEX-6P-1 vector at BamHI/XhoI sites to generate pGEX/ScAtg18. All inserts were sequenced at the Automatic DNA Sequencing Facility of CCMB to ensure that they were free of undesired mutations. pGEX/PfAtg18syn, pGEX/PIPm, pGEX/ALCAm and pGEX/ScAtg18 were transformed into BL21(DE3) *E. coli* cells, which would express the recombinant proteins as GST-fusions. pGEX-6P-1 was also transformed into BL21(DE3) *E. coli* cells to produce recombinant GST. IPTG induced cell pellets of all the five expression clones (pGEX/PfAtg18syn, pGEX/PIPm, pGEX/ALCAm, pGEX/ScAtg18 and pGEX6P1) were resuspended in lysis buffer (PBS with 1 mM DTT and 1 mg/ml lysozyme; at 5 ml/g weight of the pellet), incubated for 30 min at 4°C, and sonicated (5 secs pulses at 20% amplitude for 5-30 min depending on the sample volume). The lysate was centrifuged and the supernatant was incubated with Protino® Glutathione Agarose 4B for 30 min at 4°C. The resin was washed three times with PBS and bound proteins were eluted (50 mM Tris, 20 mM GSH, pH 7.5). Elution fractions were assessed for purity by SDS-PAGE, fractions containing pure proteins were pooled and concentrated (Amicon Ultra centricons: 10 kDa cut off for GST, 50 kDa cut off for GST/PfAtg18syn, GST/PIPm, GST/ALCAm and GST/ScAtg18) with simultaneous buffer exchange to 20 mM Tris-Cl, 50 mM NaCl, pH 8.0 at 4°C. The proteins were quantitated using BCA and used for various assays.

PfMDR1 is a 1419 amino acid residue long multi-pass integral membrane transporter. It is predicted to contain 11 transmembrane helices, a large inside domain and a large cytoplasmic domain. The PfMDR1 coding regions corresponding to cytoplasmic (1054-1419 aa) and inside (341-788 aa) domains were amplified from *P. falciparum* genomic DNA using MDR1cyt-F/MDR1cyt-R and MDR1in-F/MDR1in-R primers, and cloned into the pGEX-6P-1 vector at BamHI-XhoI site to generate pGEX/MDR1cyt and pGEX/MDR1in plasmids, respectively. These plasmids were digested with KpnI-XhoI and ligated with a similarly digested 2× c-Myc-coding sequence to obtain pGEX/MDR1cyt-myc and pGEX/MDR1in-myc plasmids, which were transformed into BL21(DE3) *E. coli* cells for expression of recombinant cytoplasmic (GST/MDRCD) and inside (GST/MDRID) proteins, respectively. Recombinant GST/MDRCD and GST/MDRID domains were purified as described above for GST-fusion proteins.

### Generation of anti-PfAtg18 antibodies

Two 4-6 weeks old male Wistar rats were immunized intraperitoneally with recombinant Trx/His-PfAtg18ct in complete (day 0) or incomplete (days 15, 30, 60, 90 and 120) Freund’s adjuvant. Sera were collected (on days 75, 105, 135 and 140), assessed for reactivity with the recombinant protein, and the day 140 serum was purified against the lysate of pET32a-transformed BL21-CodonPlus(DE3)-RIL cells to remove cross-reactive antibodies to *E. coli* proteins as has been described earlier [5]. The purified antibodies were stored in 50% glycerol with 0.01% sodium azide at −30°C.

### Western blotting

For assessing reactivity and specificity of purified anti-Atg18 antibodies, the mock-transformed *E. coli* lysate, the total lysate of induced PfAtg18ct-expressing *E. coli*, purified Trx/His-PfAtg18ct, lysates of wild type and recombinant D10 parasites, and RBC lysate were resolved on 12% SDS-PAGE gel. The proteins were transferred onto the Immobilon-P membrane, which was incubated with blocking buffer (3% skim milk in TBST), followed by incubation with rat anti-Atg18 (at 1/5000 dilution in blocking buffer) or mouse anti-β actin (1/500 dilution in blocking buffer) antibodies. The membrane was washed with blocking buffer, incubated with secondary antibodies (goat anti-Rat IgG-HRP or goat anti-Mouse IgG-HRP at 1/20,000 dilution in blocking buffer), washed again with TBST, and the signal was developed using the SuperSignal Chemiluminescent substrates. The β-actin band intensities were used as a loading control for samples in all western blots.

To determine expression of native PfAtg18 in different asexual stages of *P. falciparum* 3D7, parasite pellets of ring, early trophozoite, late trophozoite and schizont stages were lysed with SDS-PAGE sample buffer (1× sample buffer contains 20% v/v glycerol, 4% w/v SDS, 0.25% bromophenol blue, 0.1% v/v β-ME and 0.25 M Tris-Cl, pH 6.8), centrifuged at 23755g for 20 min, the supernatants were separated and equal amounts of supernatant samples were processed for western blotting using anti-Atg18 or mouse anti-β-actin antibodies, followed by appropriate secondary antibodies as described above.

### Construction of transfection plasmids

The complete coding sequence of PfAtg18 was amplified from the p2.1-ATG18 plasmid using Atg18expF/PfAtg18-Rep primers, and cloned into the pGT-GFPbsc plasmid at BamHI/XhoI sites to generate pGT-GFP/PfAtg18 construct. The GFP/PfAtg18 insert was subcloned at BglII/XhoI sites in pPfCENv3 and pSTCII-GFP plasmids to obtain pCENv3-GFP/PfAtg18 and pSTCII-GFP/PfAtg18 plasmids, respectively. pPfCENv3 and pSTCII-GFP plasmids have been described previously [57, 58]. The *P. vivax* Atg18 (PvAtg18) coding sequence was amplified from *P. vivax* genomic DNA using PvA18 F/PvA18 R primers, cloned into pGT-GFPbsc plasmid at BamHI/XhoI sites to generate pGT-GFP/PvAtg18 plasmid, from which the GFP/PvAtg18 insert was subcloned into pSTCII-GFP plasmid at BglII/XhoI sites to obtain pSTCII-GFP/PvAtg18 transfection plasmid. The *P. knowlesi* Atg18 (PkAtg18) was amplified using PkA18-F/PkA18-R primers from the *P. knowlesi* genomic DNA, cloned into the pGT-GFPbsc vector at BamHI/XhoI sites to generate pGT-GFP/PkAtg18, which was further subcloned into pSTCII-GFP vector at BglII/XhoI sites to obtain pSTCII-GFP/PkAtg18 transfection plasmid. The PvAtg18 and PkAtg18 genes contain a single intron, which would be spliced from the transcript by the parasite splicing machinery.

PIPm and ALCAm mutants of PfAtg18 were generated by recombination PCR method using primers containing the desired mutations (FRRG to FAAG in PIPm and WLCL to ALCA in ALCAm) [59]. PIPm was amplified as two fragments from the pSTCII-GFP/PfAtg18 plasmid using GFPseqF/PfA18-mPIP-R and PfA18-mPIP-F/PfAtg18-Rep primers. The two fragments were recombined using GFPseqF/PfAtg18-Rep primers. ALCAm was amplified as two fragments from the pSTCII-GFP/PfAtg18 plasmid using the primer sets GFPseqF/PfA18-mLIR-R and PfA18-mLIR-F/PfAtg18-Rep. The two fragments were recombined using GFPseqF/PfAtg18-Rep primers. PIPm and ALCAm fragments were digested with NcoI/XhoI and cloned into similarly digested pSTCII-GFP to obtain pSTCII-GFP/PIPm and pSTCII-GFP/ALCAm transfection plasmids. GFP/PIPm and GFP/ALCAm fragments were excised from the respective plasmids with BglII/XhoI and ligated into the similarly digested pPfCENv3 vector to obtain pCENv3-GFP/PIPm and pCENv3-GFP/ALCAm transfection constructs, respectively. ScAtg18 was excised from the pGT-GFP/ScAtg18BmBgm plasmid and cloned into the pPfCENv3 plasmid at BglII/XhoI sites to obtain the pCENv3-GFP/ScAtg18 transfection plasmid.

For construction of *P. berghei* Atg18 (PbAtg18) knockout plasmid, 5’UTR (flank1) and 3’UTR (flank2) of PbAtg18 were amplified from *P. berghei* genomic DNA using PbA18KFL1-F1/PbA18KFL1-R1 and PbA18KFL2-F/PbA18KFL2-R1 primers, respectively. The flank1 and flank2 were sequentially cloned into the HB-DJ1KO plasmid [60] at NotI/KpnI and AvrII/KasI, respectively, to generate HBA18KO transfection plasmid. For construction of a PbAtg18 knock-in plasmid, the GFP insert was excised from the pGT-GFPbsc plasmid and subcloned into the HBA18KO plasmid at KpnI/XhoI to obtain HBA18GFP plasmid. The PbAtg18 coding region with 5’UTR was amplified from *P. berghei* genomic DNA using PbA18KFL1-F1/PbA18Rep-R primers and cloned into the HBA18GFP plasmid at NotI/KpnI sites, replacing the 5’UTR, to construct HBA18/GFPki transfection plasmid. For construction of a PbAtg18 knock-down plasmid, the mutant *E. coli* DHFR coding sequence with HA-tag (cDD_HA_) was amplified from pPM2GDBvm plasmid (a kind gift from Dr. Praveen Balabaskaran Nina) using cDD-F/cDD-R primers and cloned into the pGT-GFPbsc plasmid at KpnI/XhoI sites to obtain pGT-cDD_HA_ plasmid. The cDD_HA_ was excised from the pGT-cDD_HA_ plasmid and subcloned into the HBA18KO plasmid at KpnI/XhoI sites to obtain HBA18cDD_HA_ plasmid. The PbAtg18 coding region was amplified from *P. berghei* genomic DNA using PbA18Rep-F2/PbA18Rep-R primers and cloned into the HBA18cDD_HA_ plasmid at NotI/KpnI sites, replacing the 5’UTR flank1, to obtain HBA18/cDD_HA_kd transfection plasmid. The HBA18KO, HBA18/GFPki and HBA18/cDD_HA_kd plasmids were digested with EcoRV to obtain linear transfection constructs, which were used for transfection of *P. berghei* ANKA.

The PfCRT coding region was amplified from *P. falciparum* cDNA using PfCRT-Fepi/PfCRT-Repi primers and cloned into the pGEM-T easy vector to generate pGEM-PfCRT plasmid. The mCherry coding region was amplified from the pmCherry-N1 (Clontech) plasmid using mCher-F/mCher-R primers and cloned into the pGT-GFPbsc plasmid at BglII/XhoI sites, replacing the GFP region, to obtain pGT-mCherry. The PfCRT insert was excised from the pGEM-PfCRT plasmid with BglII/KpnI and subcloned into the similarly digested pGT-mCherry plasmid to generate pGT-PfCRT/mCherry. The HB-DJ1KO plasmid was modified to express PfCRT/mCherry under 5’UTR and 3’UTR of *P. berghei* DNA damage inducible-1 protein (PbDdi1). The PbDdi1 5’UTR and 3’UTR were amplified from the *P. berghei* genomic DNA using PbDdi1-Fl1F/PbDdi1-Fl1R and PbDdi1-Fl2F/PbDdi1-Fl2R primers, respectively. The Ddi1 5’UTR was digested with NotI/KpnI and the Ddi1 3’UTR was digested with AvrII/KasI, and sequentially cloned into the similarly digested HB-DJ1KO plasmid to obtain HB-Ddi plasmid. The PfCRT/mCherry fragment was excised from the pGT-PfCRT/mCherry plasmid with KpnI/XhoI and cloned into the similarly digested HB-Ddi plasmid to obtain HB-PfCRT/mCherry transfection plasmid.

The coding regions and flanks in all the transfection plasmids were sequenced to ensure that they were free of undesired mutations, and the presence of different regions was confirmed by digestion with region-specific restriction enzymes. The transfection constructs were purified using the NucleoBond® Xtra Midi plasmid DNA purification kit.

### Transfection of *P. falciparum*

Early ring stage-infected RBCs from freshly thawed cultures of *P. falciparum* 3D7 and D10 strains were transfected with 50-100 µg of desired transfection plasmid DNAs (pCENv3-GFP/PfAtg18, pCENv3-GFP/PIPm, pCENv3-GFP/ALCAm, pCENv3-GFP/ScAtg18, HB-PfCRT/mCherry) as has been described earlier [61, 62]. The selection of transfected parasites was started with appropriate drugs (blasticidin at1 µg/ml for pCENv3-based plasmids and WR99210 at 1 nM for HB-based plasmids) from day 2-12, followed by no drug till day18, and then reapplication of the respective drug till the emergence of resistant parasites. Recombinant parasites were usually maintained in the presence of blasticidin, which was withdrawn during experiments. For co-transfection, the pCENv3-GFP/PfAtg18-transfected *P. falciparum* D10 parasites were cultured in the absence of blasticidin for 2 cycles. The ring stage-infected RBCs were transfected with HB-PfCRT/mCherry plasmid, and subjected to selection (0.5 µg/ml blasticidin and 1 nM WR99210) from day 2-12, and thereafter maintained in the presence of 1 nM WR99210 till the emergence of resistant parasites.

### Transfection of *P. berghei*

*P. berghei* ANKA was maintained in BALB/c mice as described in the parasite culture section. The infected mice were euthanized at 6-8% parasitemia and the blood was collected in Alsever’s solution by cardiac puncture. The trophozoite stage parasites were purified on a 65% Histodenz gradient at 360g with 0 deceleration using a swinging-bucket rotor, and cultured in RPMI 1640-FBS medium (supplemented with 2 g/litre sodium bicarbonate, 2 g/litre glucose, 25 µg/ml gentamicin, 300 mg/litre glutamine and 20% FBS) for 12-15 hrs at 35°C with shaking at 50 rpm. The culture was centrifuged at 1398g for 5 min and the parasite pellet, which mostly contained mature schizonts, was resuspended in 100 µl Nucleofector solution (Lonza) containing ∼5 µg of desired circular (pSTCII-GFP/PfAtg18, pSTCII-GFP/PvAtg18, pSTCII-GFP/PkAtg18, pSTCII-GFP/PIPm, pSTCII-GFP/ALCAm) or linear (HBA18KO, HBA18/GFPki and HBA18/cDD_HA_kd) transfection plasmids. The content was electroporated using Amaxa Nucleofector device as has been described previously [63]. 120 µl of RPMI 1640-FBS medium was added to the cuvette, and the entire sample was injected intravenously into a naive mouse. The mouse was given pyrimethamine in drinking water (70 µg/ml) for 7 days for selection of transfected parasites. For transfections with HBA18/cDD_HA_kd, mice were given water with trimethoprim (400 µg/ml) for 2-3 hrs before the infection, and then given water containing trimethoprim+pyrimethamine unless otherwise mentioned. Pure clonal lines of HBA18/cDD_HA_kd transfected parasites were obtained by the dilution cloning method, which involved intravenous infection of 10 mice, each with 0.5 parasite. The parasites were harvested for various analyses as mentioned elsewhere.

### Localization of wild type and mutant Atg18 proteins in parasites

For localization of PfAtg18 and its mutants, GFP/PfAtg18-, GFP/PIPm- and GFP/ALCAm-expressing *P. falciparum* and *P. berghei* parasites were processed for live cell imaging. 100 µl of parasite cultures (∼5% parasitemia) were washed with PBS, stained with Hoechst (10 µg/µl in PBS; Invitrogen), layered on poly L-Lysine coated slides, unbound cells were washed off with PBS, coverslips were mounted over the slides, and the cells were observed under the 100× objective of a ZEISS Apotome microscope. Images were taken (Zeiss AxioCam HRm) and analysed with the Axiovision software. For Z-section images, GFP/PfAtg18-expressing *P. falciparum* parasites were washed with PBS, immobilized on a poly L-Lysine coated slide, and fixed (3% paraformaldehyde and 0.01% glutaraldehyde). The cells were permeabilized (0.5% Triton X-100 in PBS), blocked (3% BSA in PBS), and incubated with DAPI (10 µg/ml in PBS; Invitrogen). The slides were mounted with ProLong Gold anti-fade (Thermo Fisher Scientific), images were captured using the Leica TCS SP8 confocal laser scanning microscope and edited using the Leica Application Suite software.

The localization of GFP/PfAtg18, GFP/PIPm, GFP/ALCAm, GFP/PvAtg18 and GFP/PkAtg18 was assessed in *P. berghei* trophozoites and schizonts. For localization in trophozoites, 10-20 µl blood was collected in Alsever’s solution from the tail snips of infected-mice at 4-5% parasitemia, washed with PBS and processed for live cell fluorescence microscopy as described for GFP/PfAtg18-expressing *P. falciparum* parasites. For localization in schizonts, 50-100 µl blood was collected by retro-orbital bleeding and cultured in 10 ml RPMI 1640-FBS medium to obtain schizont stage as described in the transfection of *P. berghei* section. The culture was processed for live cell fluorescence microscopy as described for GFP/PfAtg18-expressing *P. falciparum* parasites.

Asynchronous cultures of GFP/PfAtg18-, GFP/PIPm-, GFP/ALCAm-, GFP/PvAtg18-, and GFP/PkAtg18-expressing *P. falciparum* and *P. berghei* were harvested at 10-15% parasitemia by saponin lysis. The pellets were processed for western blot using mouse anti-GFP (at 1/500 dilution in blocking buffer) or mouse anti-β-actin antibodies, followed by detection with appropriate secondary antibodies as described in the Western blotting section.

GFP/ScAtg18-expressing *P. falciparum* parasites were processed for live cell microscopy and western blotting as described for GFP/PfAtg18-expressing *P. falciparum* parasites.

### Colocalization of PfAtg18

For colocalization with lysotracker, GFP/PfAtg18-expressing *P. falciparum* trophozoites were incubated with 200 nM lysotracker (Invitrogen) for 1 hour, washed with PBS, stained with Hoechst, and processed for live cell fluorescence microscopy. For colocalization with PfCRT, *P. falciparum* trophozoites transfected with pCENv3-GFP/PfAtg18 and HB-PfCRT/mCherry plasmids were processed for live cell fluorescence microscopy as described for GFP/PfAtg18-expressing *P. falciparum* parasites.

### PI3P binding assay

20 µl slurry of PI(3)P Beads (Echelon Biosciences) was washed and equilibrated with 500 µl of cold binding buffer (50 mM Tris, pH 7.5, 150-200 mM NaCl, 0.25% NP-40). The beads were incubated with 2-3 µg of recombinant GST/PfAtg18syn, GST/PIPm, GST/ALCAm, GST/ScAtg18 and GST proteins for 2 hrs at room temperature with gentle shaking. The beads were washed four times with the binding buffer and bound proteins were eluted by boiling the beads in 120 µl of 1× SDS-PAGE sample buffer for 10 min. Aliquots of the eluates, input, flow-through and washes were processed for western blotting using mouse anti-GST (at 1/1000 dilution; Invitrogen), followed by goat anti-Mouse IgG-HRP antibodies as described in the western blotting section.

### Protein-protein overlay assay

For assessing interaction between PfAtg18 and PfMDR1, different amounts of purified recombinant GST, GST/PfAtg18syn and GST/ScAtg18 were immobilized on nitrocellulose membrane, the membrane was blocked (3% BSA in PBST), and then overlaid with purified recombinant GST/MDRCD or GST/MDR1ID (at 2.5 µg/ml in blocking buffer) for 12-13 hrs at 4°C. The membrane was washed with blocking buffer, incubated with mouse anti-myc antibodies, followed by HRP-conjugated goat anti-Mouse IgG as described in the western blotting section. Binding of GST/MDRCD with GST/PfAtg18syn, GST/PIPm and GST/ALCAm was compared by ELISA. Recombinant proteins (GST/PfAtg18syn, GST/PIPm, GST/ALCAm and GST) were coated to the wells of an ELISA plate (19.5 nM protein/well in 0.2 M bicarbonate buffer, pH 9.2) at 4°C for 8-10 hrs. The wells were washed with PBST (PBS with 0.1% Tween 20), and then filled with blocking buffer (3% BSA in PBST) for 2 hrs at room temperature. The blocking buffer was discarded, serial 2-fold dilutions of GST/MDRCD (in 200 µl of PBST) were added to the wells, the plate was incubated at 4°C for 12-13 hrs. The wells were washed 5 times with PBST, each time for 5 min. Mouse anti-myc antibodies (1/1000 in blocking buffer, 200 µl/well) were added to each well, and the plate was incubated for 1 hr at room temperature. The wells were washed 4 times with PBST, each time for 5 min, followed by once with blocking buffer. HRP-conjugated goat anti-mouse IgG (1/4000 dilution in blocking buffer, 200 µl/well) was added to the wells, incubated for 45 min, and washed 5 times with PBST, each time for 5 min. The reaction was developed using TMB/H_2_O_2_ for 15 min, stopped with 1N HCl, and absorbance was measured at 450 nm using the BioTek Power wave XS2 plate reader. The assay was done in triplicates for each protein concentration. The reactivity obtained with GST was subtracted from those of GST/PfAtg18syn, GST/PIPm or GST/ALCAm for the respective GST/MDRCD concentration to adjust for background reactivity. The mean of background-adjusted reactivity was plotted against GST/MDRCD concentrations using the GraphPad Prism.

### CD spectroscopy

CD spectra for recombinant GST/PfAtg18syn and GST/ALCAm proteins (0.5 mg/ml in PBS) were recorded on a Jasco J-815 spectropolarimeter at room temperature. 0.02 cm and 1.0 cm path length quartz cuvettes were used for far-UV and near-UV CD measurements, respectively. The instrument was set on wavelength spectrum scan mode and the spectra were recorded from 190 nm to 250 nm and 250 nm to 350 nm for far-UV and near-UV CD measurements, respectively. Five spectra were averaged to increase the signal-to-noise ratio, and readings were plotted using the Origin 2020b software.

### Inhibition experiments

A variety of inhibitors and antimalarials were assessed for their effects on PfAtg18 localization. The concentrations of compounds were normalized to their IC_50_ concentrations, which were either determined in this study or taken from previous reports (Table S3). Highly synchronized GFP/PfAtg18-expressing *P. falciparum* parasites at late ring (16 hour post-synchronization) or early trophozoite stage were cultured in the presence of inhibitors (22 µM E64, 300 µM pepstatin A, 30.8 nM epoxomicin, 0.48 µM pristimerin, 95 µM LY294002, 52.4 µM orlistat; all at 4x IC_50_ concentration except E64 at 3x IC_50_ concentration) or antimalarials (236.4 nM artemisinin, 120 nM chloroquine, 81.6 nM amodiaquine, 124.8 nM mefloquine, 21.6 nM halofantrine, 532 nM quinine; all at 4x IC_50_ concentration) for 8 hours. For controls, parasites were grown with DMSO (0.06% v/v) under identical conditions. The parasites were washed with PBS and processed for localization of PfAtg18 using live cell imaging as described for GFP/PfAtg18-expressing *P. falciparum* parasites. The effect of treatment on PfAtg18 localization was scored by observing at least 200 cells, and experiments were repeated at least three times for each inhibitor. For scoring the effect of chloroquine and amodiaquine at different concentrations, treatment was carried out for 4 hrs. To assess if LY294002 exerts a dose-dependent effect on PfAtg18 localization, GFP/PfAtg18-expressing *P. falciparum* parasites at late ring stage (16 hour post-synchronization) were cultured in the presence of different concentrations of LY294002 (1× IC_50_ to 5× IC_50_, where 1× IC_50_ = 23.75 μM) or DMSO (0.45% v/v) for 8 hrs. The parasites were processed for live cell imaging as described earlier for GFP/PfAtg18-expressing *P. falciparum* parasites; at least 100 cells were observed for PfAtg18 localization, and the experiment was repeated three times. To assess the effect of inhibitors or antimalarials on PfAtg18 protein level, 20-25 ml cultures (at 10-12% parasitemia) of the control and treated parasites were harvested at the end of treatment, purified by saponin lysis and processed for western blotting with mouse anti-GFP and anti-β-actin antibodies, followed by appropriate secondary antibodies as described in the Western blotting section. For colocalization of PfAtg18 with PfAtg8, the E64-treated parasites were processed for immunofluorescence assay as described in the colocalization section. To assess the effect of endosomal transport inhibitors on PfAtg18 localization, highly synchronized GFP/PfAtg18-expressing *P. falciparum* parasites at late ring stage (16 hour post-synchronization) were cultured in the presence of bafilomycin (100 nM) or brefeldin A (20 µM) or NH_4_Cl (13.6 mM) for 8 hours, and parasites were processed for live cell imaging as described for localization of GFP/PfAtg18-expressing *P. falciparum* parasites. To assess food vacuole integrity during inhibition experiments, PfCRT/mCherry-expressing *P. falciparum* parasites were treated at 16 hour post synchronization with chloroquine (120 nM), E64 (22 μM) or DMSO (0.08% v/v) for 8 hours. The parasites were processed for localization of PfCRT/mCherry using live cell imaging as described for GFP/PfAtg18-expressing *P. falciparum* parasites, and the effect of treatment was scored by observing at least 150-200 cells.

### Analysis of *P. berghei* Atg18 knock-in and knock-down parasites

Mixed population of *P. berghei* Atg18/GFP knock-in (PbAtg18-KI) and a clonal line of *P. berghei* Atg18/cDD_HA_ knock-down (PbAtg18-KD) were analysed for confirmation of integration of the transfection cassettes into the genome, localization and expression of Atg18/GFP or Atg18/cDD_HA_ proteins. 6-8 weeks old BALB/c mice were infected with frozen stocks of wild type *P. berghei* ANKA, PbAtg18-KI or PbAtg18-KD parasites and infection was monitored regularly by observing Giemsa stained blood smears of the infected mice. Pyrimethamine (70 µg/ml in drinking water) was given to the mice infected with PbAtg18-KI parasites once parasites were observed in the blood smear, and continued throughout the infection. The mice infected with PbAtg18-KD parasites received trimethoprim (400 µg/ml drinking water) 2-3 hours before the infection, and were maintained under trimethoprim+pyrimethamine during the entire course of infection. The mice were euthanized at 10-15% parasitemia, parasites were purified, and the parasite pellets were processed for isolation of genomic DNA and western blotting as described in the parasite culture section. To confirm integration of transfection cassettes, genomic DNAs of wild type *P. berghei* ANKA, PbAtg18-KI and PbAtg18-KD parasites were used in PCR reactions containing primers specific for the wild type PbAtg18 gene (A18Con-5UF/A18Con-3UR), 5’-integration locus (PbAtg18-KD: A18Con-5UF/PvAc Con-R; Atg18-KI: A18Con-5UF/GFP seqR), 3’-integration locus (Hrp2-SeqF/A18Con-3UR), and the control MSP1 gene (PbMsp1-Fl1-F/PbMsp1-Fl1Rgpi). The PCR reactions were run on 0.8% agarose gel, and visualized with ethidium bromide.

PbAtg18-KI parasites were processed for localization of PbAtg18/GFP using live cell fluorescence microscopy as described for GFP/PfAtg18-expressing *P. berghei* parasites. The expression of PbAtg18/GFP in PbAtg18-KI parasites was determined using rabbit anti-GFP antibodies (1/1000 dilution in blocking buffer) as described in the Western blotting section. PbAtg18-KD parasites were assessed for expression of PbAtg18/cDD_HA_ by western blotting using rabbit anti-HA (at 1/2000 dilution in blocking buffer), followed by HRP-conjugated goat anti-rabbit antibodies (at 1/10000 dilution in blocking buffer) as described in the Western blotting section. Mouse anti-β-actin antibodies were used to detect β-actin as a loading control. For localization of PbAtg18/cDD_HA_, 10-20 µl blood was collected from the tail-snip of PbAtg18-KD-infected mouse in Alsever’s solution, the cells were washed with PBS, immobilized on a poly-L-lysine slide, fixed (PBS with 3% para-formaldehyde and 0.01% glutaraldehyde), permeabilized (PBS with 0.01% Triton-X), and blocked (3% BSA in PBS). The slides were incubated with rabbit anti-HA antibodies (at 1/200 dilution in blocking buffer), followed by Alexa Fluor 488-conjugated donkey anti-Rabbit IgG (at 1/2000 dilution in blocking buffer with 10 µg/ml DAPI). The slides were washed, air dried, mounted with Prolong^TM^ Diamond antifade and observed under the 100× objective of AxioimagerZ.1 with Apotome.

For comparison of growth rates, 6 months old BALB/c mice were divided into three groups and infected intraperitoneally with wild type or PbAtg18-KD parasites (10^6^ parasites/mouse). One group was infected with wild type *P. berghei* ANKA, 2^nd^ group was infected with PbAtg18-KD and not given trimethoprim (-TMP group) during the course of infection, the 3^rd^ group was infected with PbAtg18-KD and maintained in trimethoprim (+TMP group) starting with 2-3 hours before the infection. Blood smears were regularly evaluated for parasitemia by counting at least 2000 cells, and parasitemia was plotted against days post-infection using GraphPad Prism. Growth rates of wild type *P. berghei* ANKA and PbAtg18-KD parasites were also compared in C57BL/6J mice exactly as described for BALB/c mice. The mice with >50% parasitemia were euthanized in all growth rate experiments.

To determine the effect of PbAtg18 knock-down on expression, 6 BALB/c mice were infected intraperitoneally with PbAtg18-KD parasites and given trimethoprim from 2-3 hours before the infection till the parasitemia reached to 10-15% parasitemia. Three mice were euthanized while they were still under trimethoprim and the blood was collected for isolation of parasites (+TMP parasites). Trimethoprim was withdrawn from the remaining 3 mice, and they were euthanized 24 hours later, blood was collected for isolation of parasites (-TMP parasites). The parasite pellets were processed for western blotting using rabbit anti-HA or mouse anti-β-actin antibodies, followed by appropriate secondary antibodies as described in the western blotting section.

### Immunoprecipitation of PbAtg18 and PfAtg18

Parasites were isolated from asynchronous cultures of GFP-expressing and GFP/PfAtg18-expressing *P. falciparum* at 10-15% parasitemia as described in the parasite culture section. Wild type *P. berghei* ANKA and PbAtg18-KI parasites were maintained in 6-10 weeks old naïve BALB/c mice and parasites were isolated as described in the parasite culture section. The parasite pellet was resuspended in 5x pellet volume of the lysis buffer (10 mM Tris, 150 mM NaCl, 0.5 mM EDTA, 0.5% NP-40, pH 7.5, protease inhibitor cocktail), subjected to 5 cycles of freeze-thaw and 5 passages through a 26.5 G needle. The sample was incubated in ice for 30 min, centrifuged at 25000g for 30 min at 4°C, and the supernatant was transferred into a fresh tube. The pellet was re-extracted with 3x pellet volume of the lysis buffer as described above, and the supernatant was pooled with the first supernatant. The supernatant was incubated with GFP-Trap^R^ MA (ChromoTek) antibodies; 15 µl slurry/2 mg protein for 2 hours at 4°C with gentle shaking. The beads were washed with wash buffer 1 (10 mM Tris, 150 mm NaCl, 0.5 mM EDTA, pH 7.5, protease inhibitor cocktail), followed by with wash buffer 2 (10 mM Tris, 300 mm NaCl, 0.5 mM EDTA, pH 7.5, protease inhibitor cocktail). The bound proteins were eluted by boiling the beads in 100 µl 2x SDS PAGE sample buffer for 15 min, and the eluate was processed for western blotting and mass spectrometry. Aliquots of the eluates were assessed for the presence of GFP/PfAtg18 or PbAtg18/GFP along with appropriate control samples (input, pellet post-extraction, flow through, washes, and beads after elution) by western blotting using rabbit anti-GFP antibodies as described above in the western blotting section.

### Mass spectrometry of immunoprecipitates

The immunoprecipitate eluates were run on a 12% SDS-PAGE gel till the pre-stained protein ladder completely entered into the resolving gel. The gel was stained with coomassie blue, destained, and the gel region containing the protein band was excised. The gel piece was sequentially washed with 800 μl of 50 mM ammonium bicarbonate-acetonitrile solution (7:3 v/v), 300-800 μl of 50 mM ammonium bicarbonate, and 800 μl of acetonitrile. The gel piece was vacuum dried, resuspended in 200 µl of 10 mM DTT for 45 min at 56°C, and incubated in 200 μl of 50 mM ammonium bicarbonate-55 mM iodoacetamide solution for 30 min at room temperature. The gel piece was washed twice with 700 µl of 50 mM ammonium bicarbonate, followed by 700 µl of acetonitrile for 10 min. The gel piece was vacuum dried and treated with sequencing grade trypsin (15 ng/μl in 25 mM ammonium bicarbonate, 1 mM CaCl_2_; Promega or Roche) at 37°C for 10-16 hours. The peptides were extracted with 5% formic acid-30% acetonitrile solution, the extract was vacuum dried, dissolved in 20 µl of 0.1% TFA, desalted using C18 Ziptips (Merck), and eluted with 40 µl of 50% acetonitrile-5% formic acid solution. The eluate was vacuum dried and resuspended in 11 µl of 2% formic acid. 10 µl of this sample was run on the Q Exactive HF (Thermo Fischer Scientific) to perform HCD mode fragmentation and LC-MS/MS analysis.

Raw data files were imported into the proteome discoverer v1.4 (Thermo Fischer Scientific), analysed and searched against the Uniprot databases of *P. berghei* ANKA (ID: UP000074855), *P. falciparum* (ID: UP000001450), *M. musculus* (ID: UP000000589) and *H. sapiens* (ID: UP000005640) using the HTSequest algorithm. The analysis parameters used were enzyme specificity for trypsin, maximum two missed cleavages, carbidomethylation of cysteine, oxidation of methionine, deamidation of asparagine/glutamine as variable modifications. The precursor tolerance was set to 5 ppm and fragmentation tolerance was set to 0.05 Da. A 1% peptide FDR threshold was applied. The peptide spectral matches (PSM) and peptide identification groups of proteins were validated using the Percolator algorithm in proteome discoverer, and filtered to 1% FDR. The PSM and peptide groups passing through the FDR were exported to an excel file for analysis. The protein hits from the PbAtg18-KI sample were compared with those from the wild type *P. berghei* ANKA, and the protein hits identified in the GFP/PfAtg18 sample were compared with those in the GFP sample. Common proteins were excluded, and unique proteins common to 3 biological replicates with a minimum of 1 unique peptide were considered.

## Results

### Food vacuole localization is a conserved feature of *Plasmodium* Atg18 proteins

PfAtg18 has been shown to be expressed in erythrocytic stages and localized to vesicular structures throughout the parasite, some of which also colocalized with the food vacuole and apicoplast [47, 48, 51]. To determine the expression of native PfAtg18 in asexual erythrocytic stages, we produced the C-terminal region of PfAtg18 as a Trx/His-PfAtg18ct fusion protein and generated anti-PfAtg18 antibodies (Figure 1A). The antibodies detected a single band of the predicted size of Trx/His-PfAtg18ct (39.4 kDa) in the western blot of induced bacterial lysate and also reacted with purified Trx/His-PfAtg18ct, but did not react with the mock-bacterial lysate, indicating specificity for recombinant Trx/His-PfAtg18ct (Figure 1B). The anti-PfAtg18 antibodies recognized a single band of the size of native PfAtg18 (43.5 kDa) in the western blot of the lysates of different asexual erythrocytic stage parasites, indicating PfAtg18 expression in these stages (Figure 1C).

**Figure 1.**
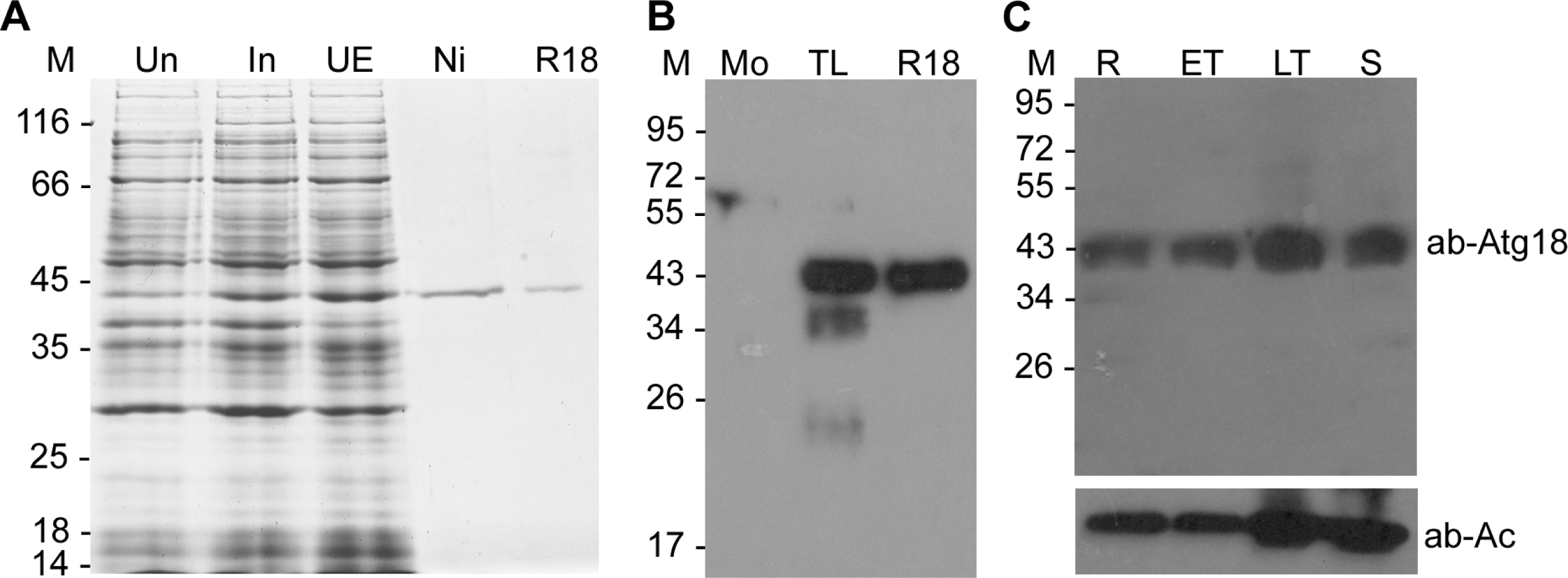
Generation of anti-Atg18 antibodies and expression of PfAtg18 in asexual erythrocytic stages. **A.** The C-terminal coding region of PfAtg18 was expressed as a Trx/His-PfAtg18ct fusion protein. The coomassie stained PAGE gel shows the lysates of unduced (Un) and IPTG-induced (In) cells, urea buffer extract of the induced cells (UE), Ni-NTA purified (Ni) and refolded Trx/His-PfAtg18ct (R18). **B.** The western blot shows reactivity of anti-PfAtg18 antibodies with the lysates of mock-transformed (Mo), total lysate of IPTG-induced cells (TL) and refolded Trx/His-PfAtg18ct (R18). **C.** A highly synchronised *P. falciparum* culture was harvested at ring (R), early trophozoite (ET), late trophozoite (LT), and schizont (S) stages. Lysates containing equal number of parasites of the indicated stages were analysed for expression of native PfAtg18 by western blotting using anti-Atg18 antibodies (ab-Atg18). β-actin was used as a loading control (ab-Ac). The protein size markers are in kDa (M).

Food vacuole, a lysosome-equivalent organelle wherein haemoglobin is degraded, has two major morphologies in *Plasmodium* species: *P. falciparum* exhibits a single large food vacuole in all asexual erythrocytic stages, whereas other *Plasmodium* species, including *P. vivax*, *P. knowlesi* and *P. berghei*, exhibit multiple vesicles most of which fuse into a single large vesicle in the schizont stage [64]. Hence, we investigated the localization of selected *Plasmodium* Atg18 proteins in *P. falciparum* and *P. berghei*, which represent the two food vacuole morphologies. For PfAtg18 localization in *P. falciparum*, it was expressed as a C-terminal fusion of GFP (GFP/PfAtg18) using an episomally maintained plasmid. The western blot of GFP/PfAtg18-expressing *P. falciparum* parasites with anti-Atg18 antibodies detected a band of the size of GFP/PfAtg18 (74.3 kDa) in addition to the native PfAtg18 (43.5 kDa), which was also detected in the lysate of wild type parasites (Figure 2A). Anti-GFP antibodies detected a single band of the size of GFP/PfAtg18 in the lysate of GFP/PfAtg18-expressing *P. falciparum* parasites, and it did not react with the lysate of wild type parasites (Figure 2B), confirming the expression of GFP/PfAtg18 in recombinant parasites. Both the antibodies did not react with uninfected RBC lysate, whereas anti-β-actin antibodies detected β-actin in all the lysates. Live cell microscopy of different stages of GFP/PfAtg18-expressing *P. falciparum* parasites showed fluorescence as a single dot in rings, whereas trophozoites and schizonts contained fluorescence around and within the food vacuole, indicating that PfAtg18 is localized to the food vacuole (Figure 2C). PfAtg18 colocalized with the lysosome marker dye lysotracker and the *P. falciparum* chloroquine resistance transporter (PfCRT), a food vacuole membrane protein (Figure S1), which further substantiated PfAtg18 localization to the food vacuole. A deeper observation of PfAtg18 localization in trophozoites, which have more pronounced food vacuole than other erythrocytic stages, revealed that it was localized both around the food vacuole (45.4% ± 9.8 parasites) and within the food vacuole (54.6% ± 9.8 parasites) (Figure S2).

**Figure 2.**
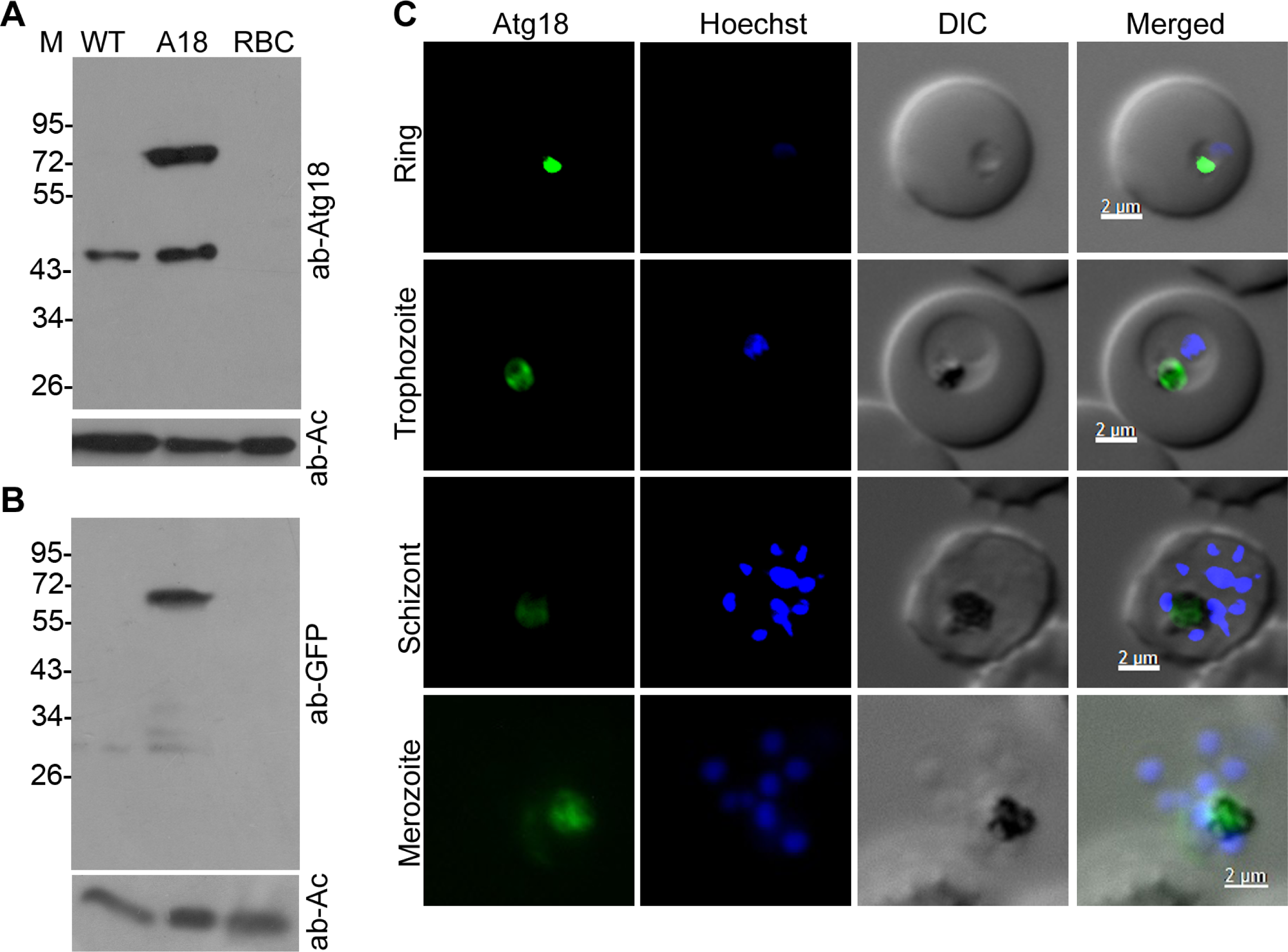
PfAtg18 localizes to the food vacuole. The GFP/PfAtg18-expressing *P. falciparum* D10 parasites were evaluated for expression (A and B) and localization (C) of GFP/PfAtg18. Western blots of the lysates of wild type parasites (WT), GFP/PfAtg18-expressing parasites (A18) and uninfected-RBCs (RBC) were probed using anti-PfAtg18 antibodies (ab-Atg18) in A and anti-GFP antibodies (ab-GFP) in B. β-actin was used as a loading control (ab-Ac). The protein bands around 43.5 kDa and 74.3 kDa correspond to native PfAtg18 and GFP/PfAtg18, respectively. The protein size markers are in kDa (M). **C**. Live GFP/PfAtg18-expressing parasites of the indicated stages were assessed for localization of GFP/PfAtg18 by fluorescence microscopy. The panels show GFP/PfAtg18 signal (Atg18), nucleic acid staining (Hoechst), bright field with RBC and parasite boundaries (DIC) and the overlap of all three images (Merged). The black substance in the DIC image is the food vacuole-resident pigment haemozoin, which serves as an in-built food vacuole marker.

For localization in *P. berghei*, GFP/PfAtg18 was expressed using an episomally maintained plasmid, and its expression was confirmed by western blot using anti-GFP antibodies (Figure S3A). These parasites showed GFP/PfAtg18 signal at multiple loci, which also contained the in-built food vacuole marker haemozoin, indicating that PfAtg18 localizes to food vacuoles, even in a heterologous parasite (Figure 3). Most of the food vacuoles present in *P. berghei* trophozoite stage merge into a single large food vacuole as the trophozoite matures into the schizont. Consistently, the majority of GFP/PfAtg18 foci merged into a large single food vacuole in schizonts. To rule out any effect of heterologous expression, we generated a *P. berghei* knock-in line (PbAtg18-KI) that expressed PbAtg18/GFP under the native promoter. PCR and western blot analysis of PbAtg18-KI parasites confirmed the replacement of wild type PbAtg18 gene with PbAtg18/GFP coding sequence and expression of PbAtg18/GFP, respectively (Figure S4). PbAtg18/GFP was predominantly associated with multiple haemozoin-containing food vacuoles in trophozoites, which coalesced into a single large food vacuole in schizonts (Figure 3). We extended localization studies to Atg18 homologs of *P. vivax*, the most prevalent human malaria parasite, and *P. knowlesi*, a zoonotic malaria parasite. The GFP-fusions of *P. vivax* Atg18 (GFP/PvAtg18) and *P. knowlesi* Atg18 (GFP/PkAtg18) were expressed in *P. berghei* using episomally maintained plasmids. Both GFP/PvAtg18 and GFP/PkAtg18 were expressed (Figure S3B) and predominantly localized to food vacuoles (Figure 4), confirming that food vacuole localization is a conserved feature of *Plasmodium* Atg18 proteins.

**Figure 3.**
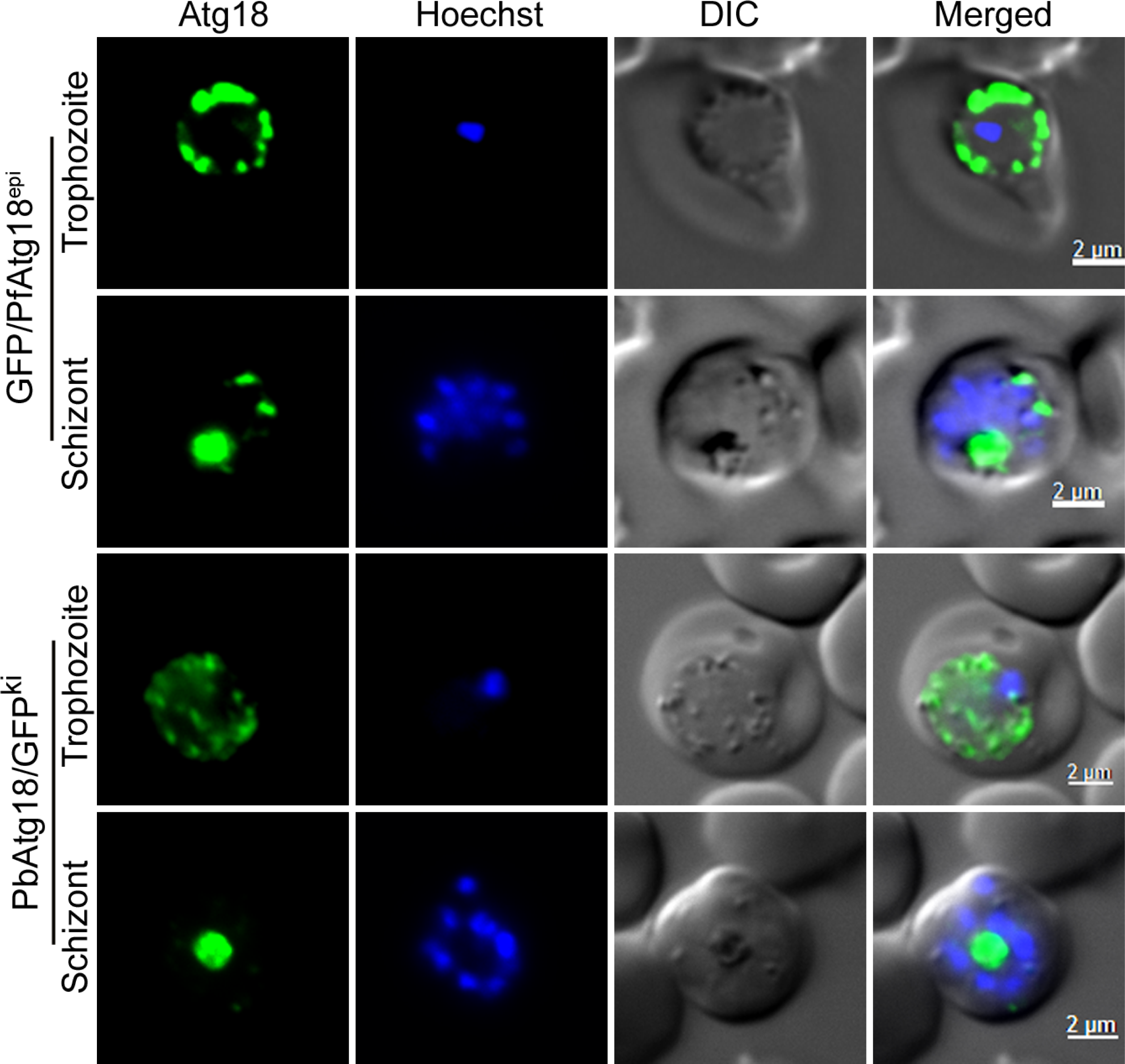
PfAtg18 and PbAtg18 show similar food vacuole localization in *P.berghei*. The GFP/PfAtg18-expressing *P. berghei* (GFP/PfAtg18^epi^) and PbAtg18/GFP knock-in *P. berghei* (PbAtg18/GFP^ki^) parasites at trophozoite and schizont stages were evaluated for Atg18 localization by live cell fluorescence microscopy. The panels are for GFP/PfAtg18 or PbAtg18/GFP signal (Atg18), nucleic acid staining (Hoechst), bright field showing the boundaries of parasite and RBC (DIC), and the overlap of all three images (Merged). Note that GFP signal is associated with multiple haemozoin-containing foci in the trophozoite stage, which fuse into one big food vacuole in the schizont stage.

**Figure 4.**
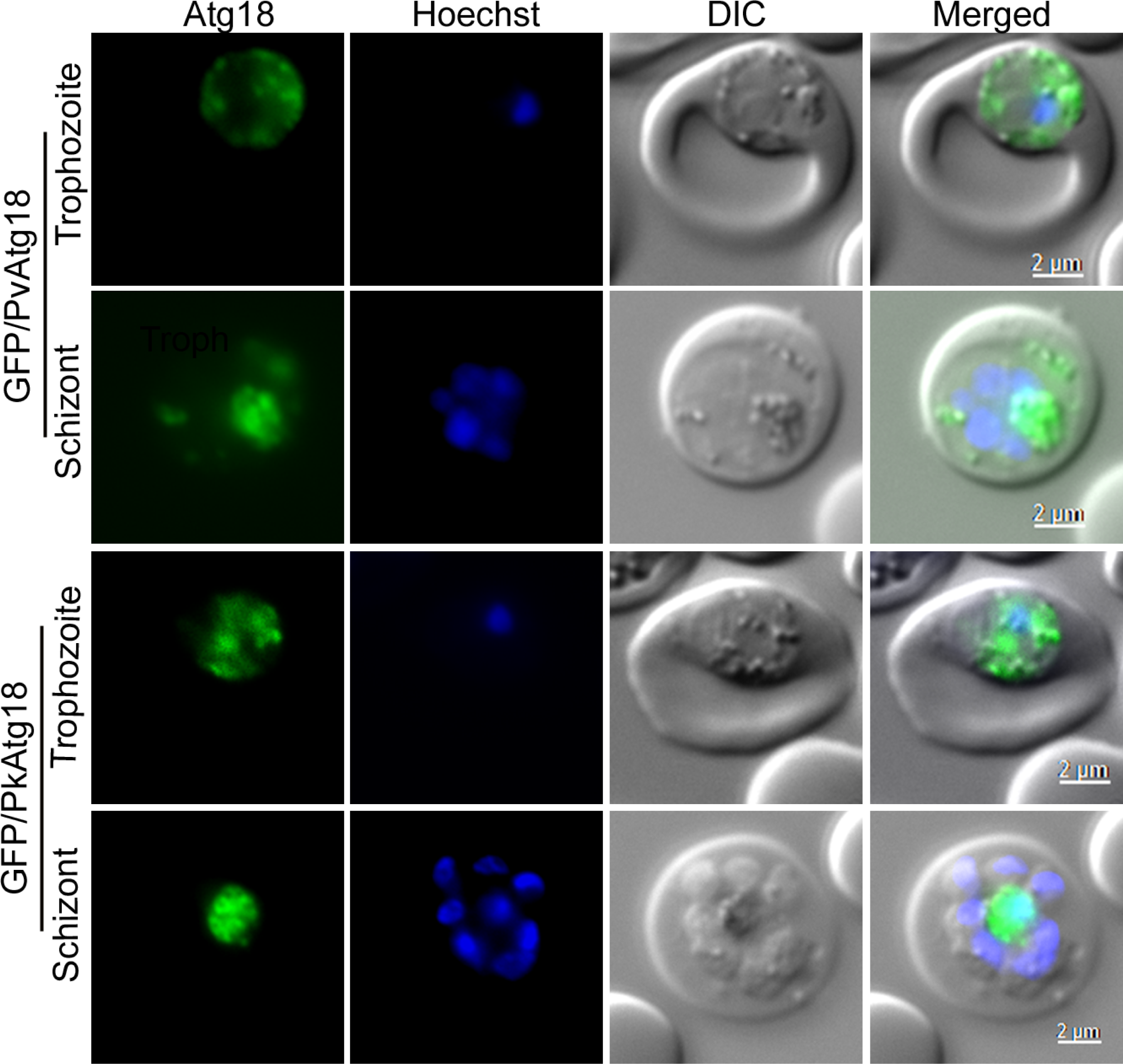
PvAtg18 and PkAtg18 localize to the food vacuoles. The GFP/PvAtg18- and GFP/PkAtg18-expressing *P. berghei* parasites at trophozoite and schizont stages were observed for localization of respective proteins by live cell fluorescence microscopy. The panels are as described in Figure 3. The GFP signal is primarily associated with multiple haemozoin-containing foci, which coalese into one big food vacuole in the schizont stage.

To investigate if PfAtg18 follows endosomal transport to the food vacuole, we evaluated its localization in GFP/PfAtg18-expressing *P. falciparum* parasites upon treatment with the inhibitors of vesicle trafficking (brefeldin A and bafilomycin) and fusion (bafilomycin and NH_4_Cl). None of the inhibitors significantly altered PfAtg18 localization (Figure S5), suggesting that it does not take endosomal trafficking route, which is in line with the absence of a signal/transmembrane sequence in PfAtg18. About 30% of the NH_4_Cl-treated parasites showed PfAtg18 around an enlarged food vacuole (Figure S5). NH_4_Cl has been shown to inhibit lysosomal degradation activity by alkalinizing the compartment [65], hence, the food vacuole enlargement could be due to accumulation of undegraded haemoglobin, as has been observed in parasites treated with E64 and pepstatin A [66–70]. Only a small fraction of NH_4_Cl-treated parasites, particularly late ring and early trophozoite stages, contained multiple PfAtg18 foci near the parasite periphery or in the cytoplasm (Figure S5), which may be due to failure of vesicles to fuse to form the food vacuole, as has also been reported earlier [71]. The PfAtg18 vesicles in NH_4_Cl-treated parasites may be part of the haemoglobin trafficking pathway, as a recent report proposed that PfAtg18 transport to the food vacuole uses haemoglobin trafficking pathway [48].

### FRRG and WLCL motifs are essential for PI3P-dependent food vacuole localization of PfAtg18

Sequence analysis using the WDSP and homology modelling on the *Pichia angusta* Atg18 structure (PDB id: 5LTD) predicted seven WD40 repeats in PfAtg18, which form a seven bladed β-propeller structure with two PI3P-binding sites, indicating that PfAtg18 is a PROPPIN family protein [18, 23], as are ScAtg18 and WIPI2 (Figure S6). The PI3P-binding sites are formed by the FRRG motif and amino acids from the 23^rd^ β strand in blade 6 (Figure S6). PfAtg18 and other *Plasmodium* Atg18 proteins also share a positionally conserved “WLCL” motif that resembles the LC3-interacting region/Atg8-interacting motif (LIR/AIM) “WxxL” in a variety of proteins, which interact with Atg8. The LIR/AIM motif is mostly preceded by negatively charged amino acids [52], whereas the “WLCL” motif is not. Nonetheless, the “WLCL” motif is moderately conserved at “WL” positions in the majority of Atg18 proteins (Figure S6) and forms the major part of 23^rd^ β strand in blade 6 that contains the 2^nd^ PI3P-binding site in yeast Hsv2 [18, 23], suggesting an important role for this motif in Atg18 function.

ScAtg18 localizes to the PAS/phagophore, vacuole and endosomes. The PAS/phagophore localization requires association of ScAtg18 with Atg2 to form the Atg18-Atg2 complex, which binds PI3P through Atg18 [18, 23, 34, 72]. Similarly, WIPI2 localizes to PI3P-rich sites on the ER, phagophore and autophagosome through interaction with PI3P [18, 23, 36, 37]. Earlier work showed that the FRRG motif of PfAtg18 is required for localization with vesicular structures in the cytoplasm and binding with PI3P [47]. Recently, PfAtg18 was shown to localize to punctate structures around the food vacuole; based on co-localization of PfAtg18 with anti-PI3P antibodies and inhibition studies using the PI3K inhibitor wortmannin, this localization was suggested to be mediated by PI3P [48]. Consistently, GFP-2xFYVE, a routinely used marker for PI3P localization studies [73], indicated abundant PI3P in the food vacuole membrane of *P. falciparum* [44]. Hence, we investigated if the food vacuole localization of PfAtg18 is mediated through its interaction with PI3P on the food vacuole membrane. We first treated GFP/PfAtg18-expressing *P. falciparum* parasites with the PI3K inhibitor LY294002 and scored the parasites for PfAtg18 localization. The treatment resulted in a dose-dependent diffuse localization of PfAtg18 throughout the parasite in a significantly larger number of parasites (18.4% ± 2.8 at 1×IC_50_ concentration to 59.8% ± 0.9 at 5×IC_50_ concentration) than in control parasites (1.6% ± 1.1) (Figure 5A), suggesting that PI3P is critical for PfAtg18 localization to the food vacuole. We next mutated the FRRG motif to FAAG, and the mutant (PIPm) was expressed as a GFP-fusion protein (GFP/PIPm) in *P*. *falciparum* and *P. berghei* (Figure S3A and S3C). The GFP/PIPm was present outside the food vacuole in the cytoplasm of all *P. falciparum* (Figure 5A) and *P. berghei* parasites (Figure S7), indicating that FRRG motif is essential for localization of PfAtg18 to the food vacuole. We also mutated the “WLCL” motif to “ALCA” and expressed the mutant (ALCAm) as a GFP-fusion protein (GFP/ALCAm) in both *P. falciparum* and *P. berghei* (Figure S3A and S3C). The GFP/ALCAm was localised outside the food vacuole in the cytoplasm of *P. falciparum* (Figure 5A) and *P. berghei* (Figure S7), indicating that “WLCL” motif is essential for localization of PfAtg18 to the food vacuole. The localization of both GFP/PIPm and GFP/ALCAm was similar to that of a cytoplasmic GFP construct (GFPcyt) (Figure 5A), confirming that FRRG and WLCL motifs are essential for the food vacuole localization of PfAtg18.

**Figure 5.**
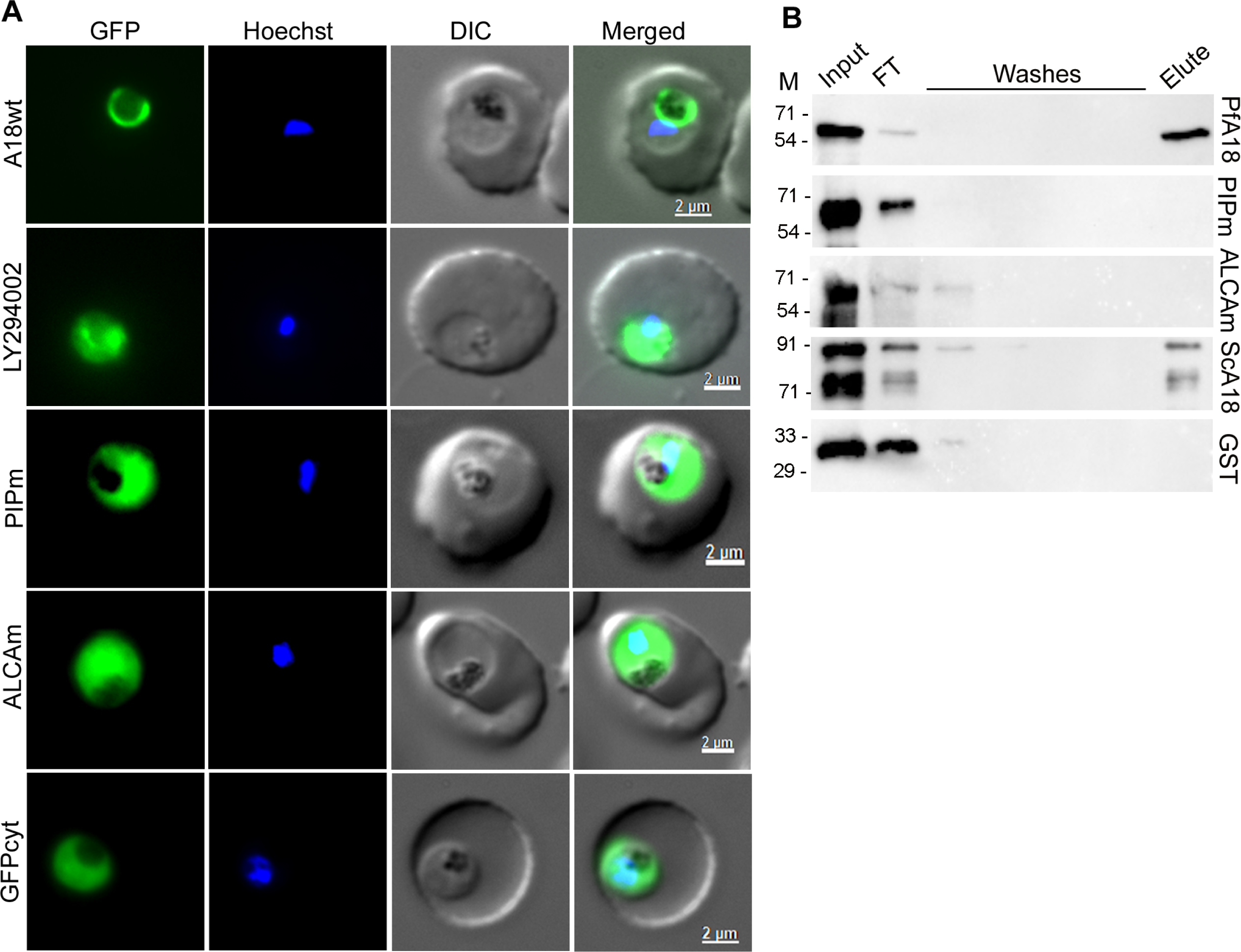
PfAtg18 localizes to the food vacuole through interaction with PI3P. **A.** GFP/PfAtg18-expressing *P. falciparum* trophozoites were treated with DMSO (A18wt) or the PI3K inhibitor (LY294002) and assessed for PfAtg18 localization by live cell fluorescence microscopy. *P. falciparum* trophozoites expressing PfAtg18 mutants (PIPm and ALCAm) were assessed for the effect of mutations on PfAtg18 localization by live cell fluorescence microscopy. Parasites expressing GFP only served as a cytosolic localization control (GFPcyt). The panels from the left represent GFP signal (GFP), nucleic acid staining (Hoechst), boundaries of parasite and RBC (DIC), and the merge of all three images (Merged). Note that the LY294002 treatment caused diffuse GFP/PfAtg18 localization all over the parasite and mutations rendered the mutant proteins outside the food vacuole in cytoplasm. **B.** Recombinant GST/PfAtg18 (PfA18), GST/PIPm (PIPm), GST/ALCAm (ALCAm), GST/ScAtg18 (ScA18) and GST were incubated for binding with PI3P-agarose beads, and the eluates (Elute) along with the input, flowthrough (FT), and washes were assessed for the presence of respective proteins by western blotting. The protein size markers are in kDa (M). The presence of signal in the eluate indicates interaction with PI3P.

To further investigate whether the loss of food vacuole localization of PfAtg18 mutants was due to their inability to interact with PI3P, recombinant GST/PfAtg18, GST/PIPm, GST/ALCAm, GST/ScAtg18 and GST were produced in BL21(DE3) cells (Figure S8), and assessed for binding with PI3P. GST/PfAtg18 bound with PI3P just like recombinant GST/ScAtg18 (Figure 5B), whereas GST/PIPm, GST/ALCAm and GST did not bind, confirming that FRRG and WLCL motifs are critical for PfAtg18 interaction with PI3P, and this interaction is essential for food vacuole localization. Since “WLCL” motif does not directly contribute to the 2^nd^ PI3P-binding site, rather, it forms the major part of 23^rd^ β strand in blade 6 that contains the 2^nd^ PI3P-binding site in yeast Hsv2, we compared CD spectra of GST/PfAtg18 and GST/ALCAm proteins. The far-UV CD spectrum of GST/ALCAm was different from that of GST/PfAtg18, whereas the near-UV CD spectra were similar, suggesting a local structural perturbation in GST/ALCAm resulting in the loss of binding to PI3P (Figure S9).

### Interaction with PI3P is not the sole requirement for food vacuole localization of PfAtg18

The “FRRG” motif is essential for targeting ScAtg18 and WIPI2 to PI3P-rich sites, and the loss of autophagic function of “FTTG” mutant of ScAtg18 can be fully restored by fusing it with the 2×FYVE domain, which specifically binds PI3P [72]. Hence, we investigated if the “FRRG” motif of ScAtg18 is sufficient to recruit it to the food vacuole of *P. falciparum*. GFP/ScAtg18 was expressed in *P. falciparum* (Figure S3D), and the parasites were assessed for localization of GFP/ScAtg18. To our surprise, GFP/ScAtg18 did not localize to the food vacuole, rather, it remained in the cytoplasm just like GFPcyt (Figure 6), indicating that interaction with PI3P alone is not sufficient and additional factors could be involved in recruitment of PfAtg18 to the food vacuole.

**Figure 6.**
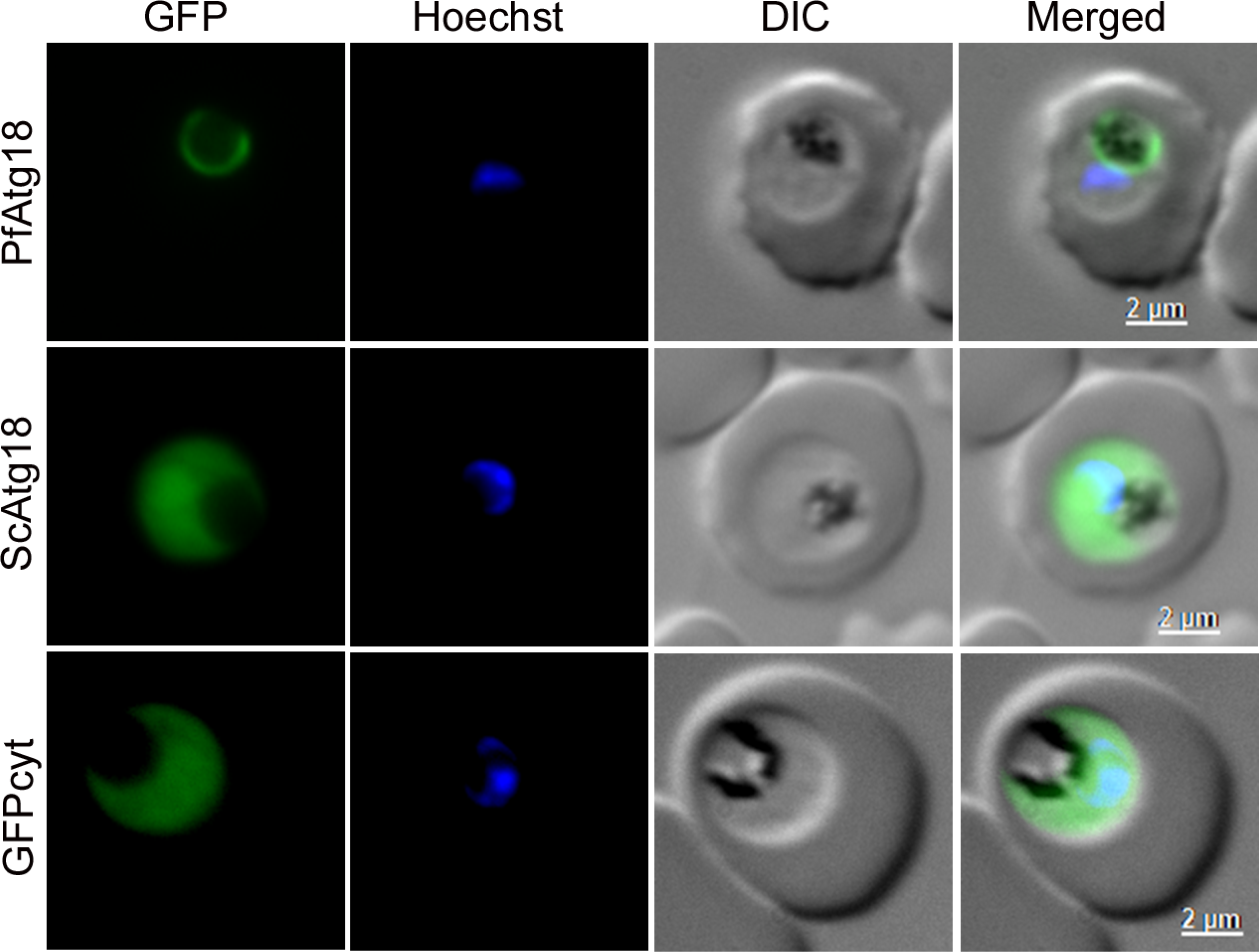
ScAtg18 does not localize to the *P. falciparum* food vacuole. The *P. falciparum* 3D7 trophozoites expressing GFP/PfAtg18 (PfAtg18), GFP/ScAtg18 (ScAtg18) and a cytoplasmic GFP control (GFPcyt) were evaluated for localization of respective proteins by live cell fluorescence microscopy. The panels are for the GFP signal (GFP), nucleic acid staining (Hoechst), bright field showing the boundaries of parasite and RBC (DIC), and the overlap of all the three images (Merged). Contrary to the food vacuole localization of GFP/PfAtg18, GFP/ScAtg18 shows diffuse cytoplasmic localization outside the food vacuole, which is similar to that of GFPcyto.

### MDR1 mediates PfAtg18 localization to the food vacuole

To identify *Plasmodium* Atg18-associated proteins, we immunoprecipitated PbAtg18/GFP and GFP/PfAtg18 (Figure S10), and the samples were subjected to mass spectrophotometry. 10 unique proteins were identified in the GFP/PfAtg18 immunoprecipitate and 37 unique proteins were identified in the PbAtg18/GFP immunoprecipitate and (Table 1, Table S1). MDR1 and DnaJ proteins were common in both the samples, suggesting association of *Plasmodium* Atg18 with MDR1 and DnaJ. The GFP/PfAtg18 immunoprecipitate also contained a HSP40 protein, a *Plasmodium* exported protein, a PHISTb protein, three ribosomal proteins, and a proteasome regulatory subunit. MDR1 is a multi-pass transmembrane protein belonging to the ABC family of transporters; it is localized on the food vacuole membrane and associated with resistance to multiple antimalarials [74–78]. MDR1 is predicted to have 11 transmembrane helices, an inside domain and a cytoplasmic domain (Figure 7A). To investigate whether PfAtg18 directly interacts with MDR1, we produced recombinant GST-tagged inside (GST/MDRID) and cytoplasmic (GST/MDRCD) domains (Figure 7B and C), and tested both the domains for interaction with GST/PfAtg18, GST/ScAtg18 and GST using dot blot protein overlay assay. GST/MDRCD, but not GST/MDRID, interacted with GST/PfAtg18 (Figure 7D). Both the recombinant MDR domains did not interact with GST and GST/ScAtg18, indicating that PfAtg18-MDR1 interaction is direct and specific. We also compared interaction of wild type PfAtg18 and its mutants with GST/MDRCD by ELISA. The PIPm and ALCAm mutants showed significantly weaker interaction with GST/MDRCD than that of wild type Atg18 (Figure S11), which substantiate PfAtg8-MDR1 interaction. Hence, MDR1, in addition to PI3P, plays a critical role in localization of PfAtg18 to the food vacuole.

**Figure. 7.**
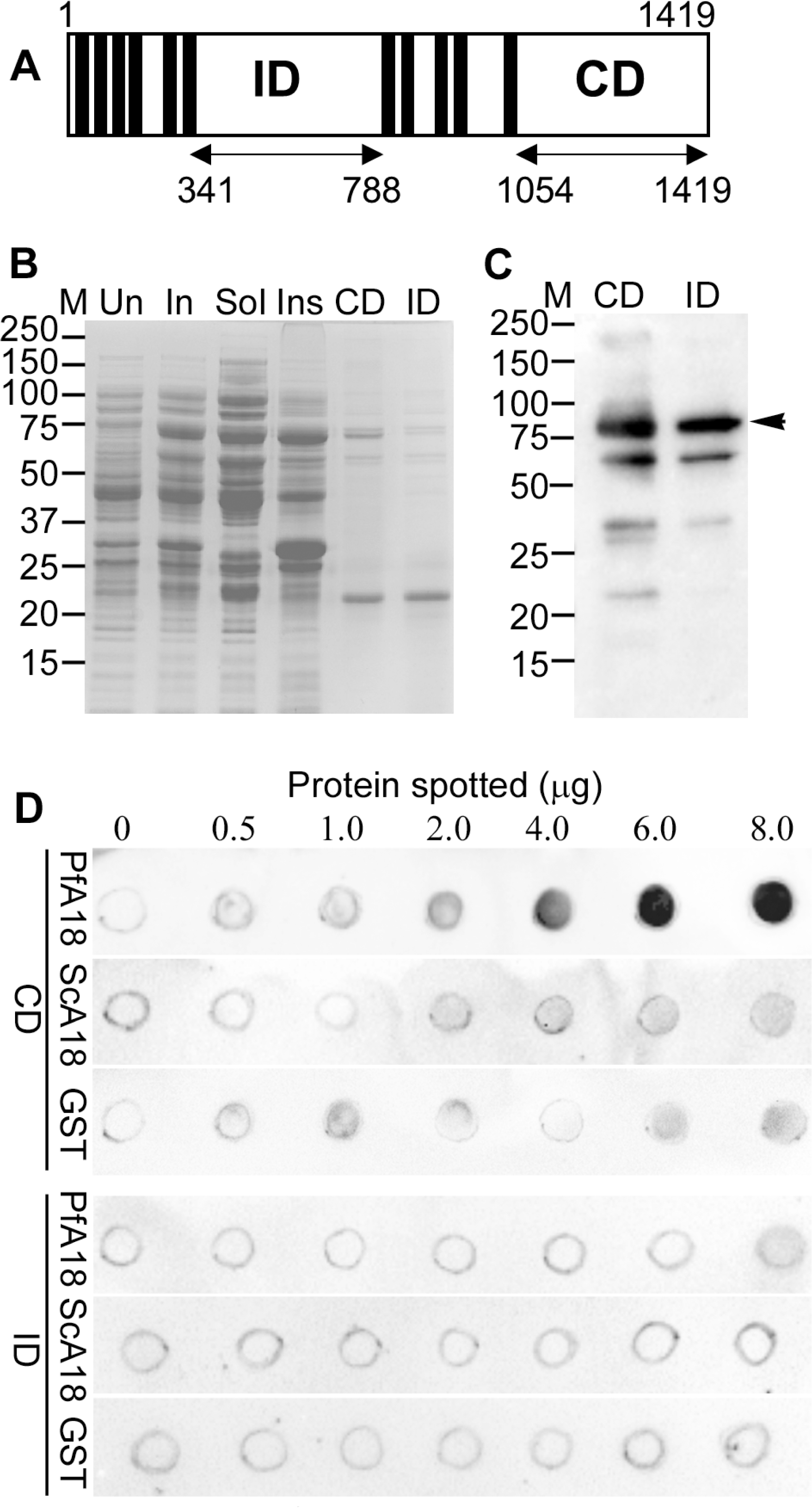
PfMDR1 interacts with PfAtg18. **A.** Schematic of the predicted domain architecture of PfMDR1 shows transmembrane helices (filled bars), the inside domain (ID) and the cytoplasmic domain (CD). GST-fusions of the cytoplasmic (GST/MDRCD) and inside (GST/MDRID) domains were expressed in *E. coli*, and enriched for fusion proteins by purification. **B.** The coomassie stained PAGE gel contains lysates of unduced (Un) and IPTG-induced (In) GST/MDRCD-expressing cells, soluble (Sol) and insoluble (Ins) fractions of the induced GST/MDRCD-expressing cells, purified GST/MDRCD (CD) and GST/MDRID (ID) proteins. **C.** The western blot shows GST/MDRCD (CD) and GST/MDRID (ID) proteins (indicated with an arrow head) in the purified samples. The protein size markers are in kDa (M). **D.** The blots containing spots of the indicated amounts of recombinant GST/PfAtg18 (PfA18), GST/ScAtg18 (ScA18) and GST were overlaid with recombinant GST/MDRCD (CD) or GST/MDRID (ID) proteins and probed with anti-GST antibodies. The blots show interaction of the PfMDR1 cytoplasmic domain with PfA18 in a dose-dependent manner.

**Table 1.** Proteins identified in the immunoprecipitate of GFP/PfAtg18-expressing *P. falciparum* parasites. The proteins listed are common in at least 3 of the 4 independent experiments, and contain at least 1 unique peptide. Shown are coverage (Cov), unique peptides (U Pep) and score (Sco) for each protein.

| Accession No | Proteins | Experiment-1 |  |  | Experiment-2 |  |  | Experiment-3 |  |  |
| --- | --- | --- | --- | --- | --- | --- | --- | --- | --- | --- |
|  |  | Cov | U<br>Pep | Sco | Cov | U<br>Pep | Sco | Cov | U<br>Pep | Sco |
| C6KTC7<br>(PF3D7_0629200) | DnaJ protein, putative | 2.4 | 1 | 2.0 | 2.4 | 1 | 4.5 | 19.5 | 7 | 36.8 |
| Q7K6A5<br>(PF3D7_0523000) | Multidrug resistance<br>protein 1 | 6.3 | 6 | 34.6 | 3.5 | 3 | 4.5 | 1.1 | 1 | 5.8 |
| Q8IL88<br>(PF14_0359) | HSP40, subfamily A,<br>putative | 7.3 | 2 | 2.3 | 13.7 | 7 | 24.6 | 2.8 | 1 | 2.8 |
| Q8IDS6<br>(PF3D7_1341200) | 60S ribosomal protein<br>L18a | 18.5 | 3 | 7.2 | 5.4 | 1 | 2.5 | 15.2 | 3 | 19.7 |
| O77395<br>(PF3D7_0316800) | 40S ribosomal protein<br>S15A, putative | 6.2 | 1 | 4.4 | 6.2 | 1 | 2.2 | 33.1 | 5 | 16.3 |
| Q8IKH8<br>(PF14_0627) | 40S ribosomal protein<br>S3 | 5.9 | 1 | 2.9 | 7.7 | 1 | 2.3 | 31.2 | 7 | 23.8 |
| Q8IEQ1<br>(PF3D7_1306400) | 26S protease<br>regulatory subunit<br>10B, putative | 3.1 | 1 | 1.9 | 5.3 | 1 | 6.2 | 27.7 | 7 | 44.2 |
| Q8I490<br>(PF3D7_0501000) | Plasmodium exported<br>protein | 8.1 | 2 | 12.2 | 4.2 | 1 | 3.1 | 13.9 | 3 | 14.9 |
| Q8I207<br>(PF3D7_0401800) | Plasmodium exported<br>protein (PHISTb) | 11.3 | 8 | 28.2 | 2.7 | 1 | 2.9 | 20.7 | 13 | 59.9 |
| Q8IJR6<br>(PF10_0126) | Autophagy-related<br>protein 18, putative | 49.0 | 18 | 184.1 | 49.2 | 19 | 235.8 | 6.6 | 2 | 7.1 |

### Food vacuole-localization of PfAtg18 is sensitive to chloroquine and amodiaquine

The food vacuole of malaria parasites is the site of some of the essential and best studied biochemical processes, including haemoglobin degradation and haemozoin formation. PfCRT and MDR1, which confer resistance to multiple antimalarials, including quinolines, are also present in the food vacuole membrane [48, 49, 75–80]. Altered distribution of Atg8 puncta and Thr38Ile mutation of PfAtg18 have been associated with resistance to chloroquine and artemisinin, respectively [12, 49–51]. However, a direct link between autophagy and drug resistance is yet to be identified. Hence, we treated GFP/PfAtg18-expressing *P. falciparum* parasites with quinolines (chloroquine, amodiaquine, mefloquine, halofantrine and quinine) and evaluated for localization of PfAtg18. Chloroquine and amodiaquine caused diffuse localization of PfAtg18 throughout the parasite in a significant number of parasites, which became more pronounced with increasing drug concentrations, whereas mefloquine, halofantrine and quinine did not affect the localization (Figures 8 and 9), indicating that the effect of chloroquine and amodiaquine on PfAtg18 localization is specific. The immunoblot of quinoline-treated parasites did not show any noticeable change in PfAtg18 levels compared to that in control parasites (Figure S12A). The treatment of GFP/PfAtg18-expressing *P. falciparum* parasites with artemisinin neither affected PfAtg18 localization (Figure 8) nor the levels (Figure S12A). ScAtg18 and PfAtg18 have been shown to participate in vacuole/food vacuole dynamics [41, 42, 48], which is the major site of haemoglobin degradation, quinoline action and source of iron for artemisinin activation. Altered food vacuole localization of PfAtg18 by amodiaquine and chloroquine could affect food vacuole-associated processes, which might modulate the effects of these drugs on the parasite.

**Figure 8.**
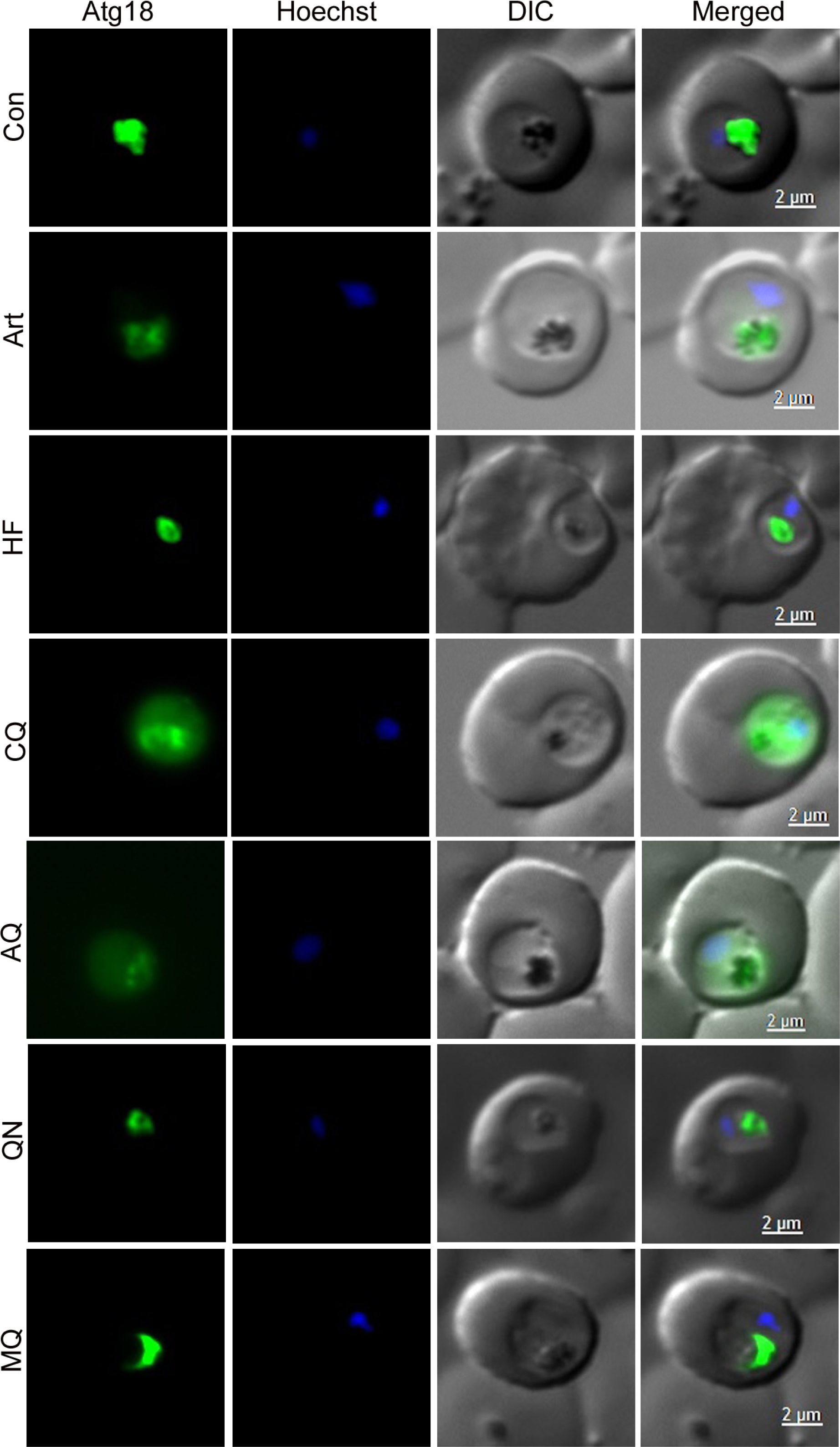
Effect of antimalarials on PfAtg18 localization. Early trophozoite stage parasites were treated with DMSO (Con) or antimalarials (artemisinin, Art; halofantrine, HF; chloroquine, CQ; amodiaquine, AQ; quinine, QN; mefloquine, MQ) for 8 hours and assessed for PfAtg18 localization by live cell fluorescence microscopy. The panels from the left are for GFP/PfAtg18 signal (Atg18), nucleic acid staining (Hoechst), bright field showing the boundaries of parasite and RBC (DIC), and the overlap of all three images (Merged). Compared with the food vacuole localization of GFP/PfAtg18 in control cells, treatment with chloroquine and amodiaquine caused diffuse localization all over the parasite.

**Figure 9.**
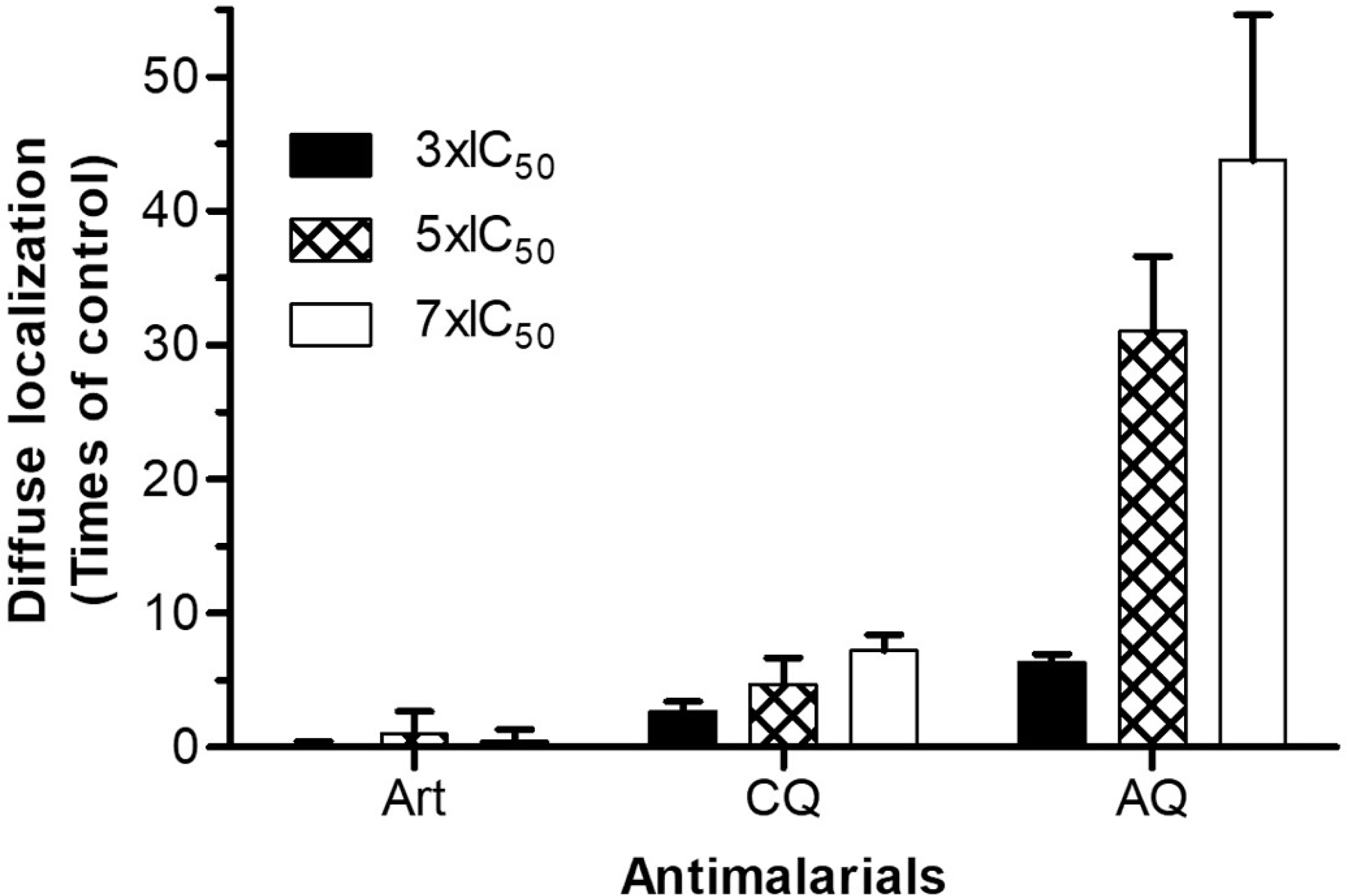
Effect of chloroquine and amodiaquine on PfAtg18 localization. GFP/PfAtg18-expressing parasites at early trophozoite stage were treated with DMSO (control) or the indicated IC_50_ concentrations of antimalarials (chloroquine, CQ; artemisinin, Art; amodiaquine, AQ) for 4 hours. The graph shows the number of parasites with diffuse localization as times of the control parasites at different IC_50_ concentrations. The data is mean with SD of three independent experiments.

### Food vacuole metabolism inhibitors affect PfAtg18 localization

Haemoglobin degradation by asexual erythrocytic stage parasites provides amino acids for parasite protein synthesis and has also been proposed to keep the infected erythrocyte osmotically stable [81, 82]. It is majorly carried out by cysteine protease falcipains and aspartic protease plasmepsins. Generic cysteine protease inhibitor E64 and aspartic protease inhibitor pepstatin A block haemoglobin degradation, resulting in the enlargement of food vacuole due to accumulation of undegraded haemoglobin [68–70], particularly in case of E64 [66–68, 83]. Although not yet reported in *Plasmodium*, lysosome is the primary site of lipid catabolism that involves transport of lipid droplets to lysosomes by lipophagy and subsequent degradation of lipid droplets by lysosomal lipases. Lipid droplets have been shown in the *Plasmodium* food vacuole, but specific food vacuole lipases are yet to be characterized.

Hence, we treated GFP/PfAtg18-expressing *P. falciparum* parasites with several inhibitors of the food vacuole-associated metabolic processes and evaluated for PfAtg18 localization. E64 and pepstatin A caused a profound ring-like localization of GFP/PfAtg18 around the food vacuole, suggesting that it is associated with the food vacuole membrane (Figure 10). The ring-like localization pattern was seen in a much larger number of parasites in case of E64 and pepstatin A than control parasites. The dot-like pattern might be of the PfAtg18 in the food vacuole lumen or due to processing of PfAtg18 on the luminal side by the food vacuole-resident proteases. A significant increase in the number of parasites with ring-like pattern upon treatment with E64 and pepstatin A supports this point. We also observed intense signal at a single site in the peri-food vacuole region, which was particularly evident in parasites treated with E64 and pepstatin A. Orlistat, an inhibitor of pancreatic lipases and fatty acid synthase [84, 85], has been shown to block the development of malaria parasites by inhibition of triacylglycerol hydrolysis [86]. A significant number of orlistat-treated parasites showed diffuse PfAtg18 localization all over the parasite, which could be due to decreased levels of fatty acids for membrane synthesis and phosphatidylinositide precursors (Figure 10). Localization of GFP/PfAtg18 in parasites-treated with pristimerin (inhibitor of monoacylglycerol lipase and proteasome) and epoxomicin (proteasome inhibitor) was nearly identical to that in control parasites (Figure 10). The immunoblot of inhibitor treated-parasites and control parasites had similar PfAtg18 levels (Figure S12B), which ruled out any effect of treatment on PfAtg18 expression level. To address whether food vacuole integrity was intact during inhibitor treatment experiments, we treated the parasites expressing PfCRTmCherry, a food vacuole marker, with E64 and chloroquine, which inhibit haemoglobin degradation and haemozoin formation in the food vacuole, respectively. In both the cases, the number of parasites showing food vacuole-associated PfCRTmCherry was comparable to DMSO control (Figure S13), indicating that food vacuole integrity was not compromised in our experiments. As observed in case of PfAtg18, the E64-treated parasites also showed an enlarged ring of PfCRTmCherry due to enlargement of the food vacuole.

**Figure 10.**
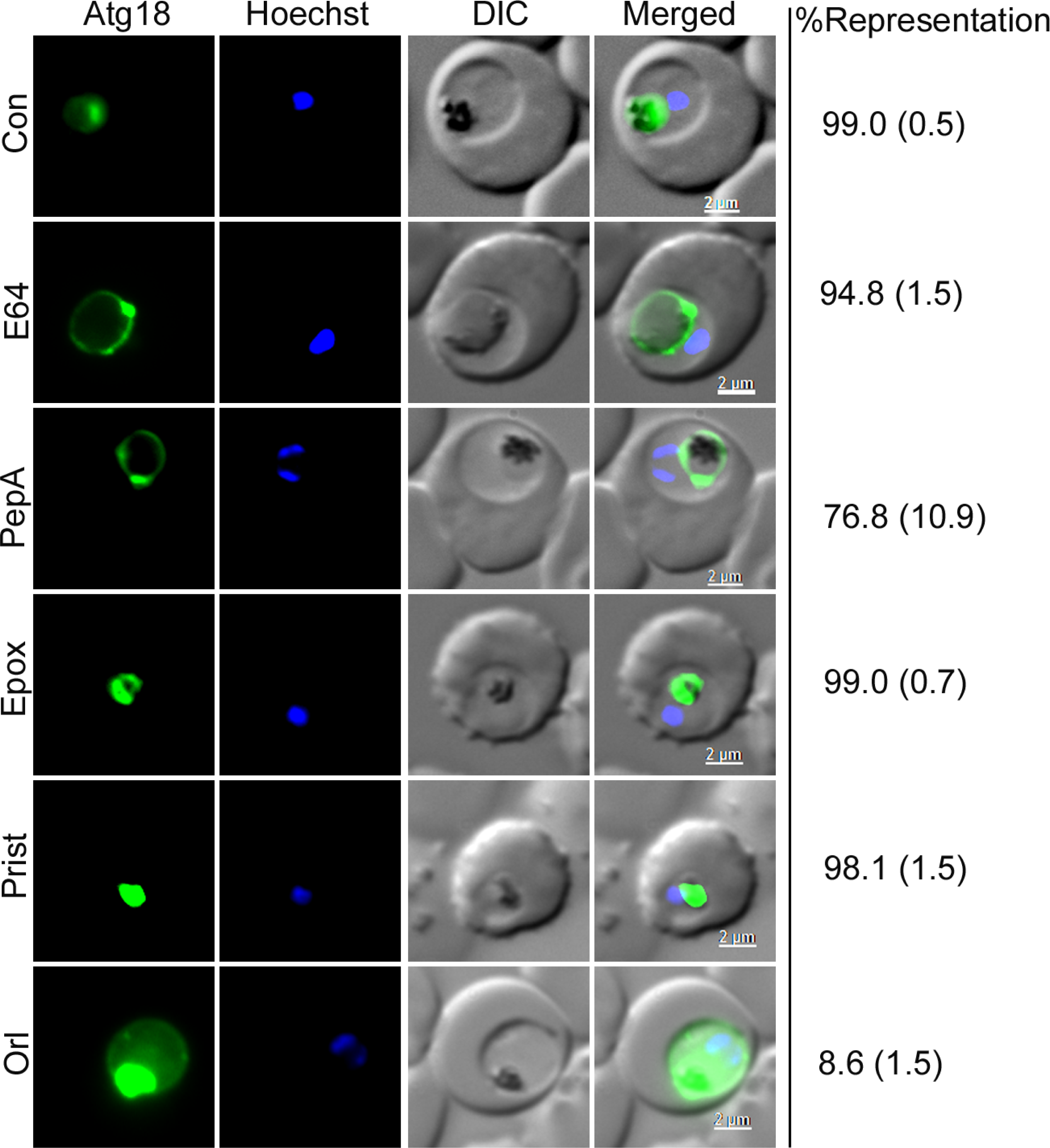
PfAtg18 localization upon treatment with the inhibitors of food vacuole-associated processes. Early trophozoite stage GFP/PfAtg18-expressing parasites were treated with DMSO (Con) or inihibitors (E64; pepstatin A, PepA; epoxomicin, Epox; pristimerin, Prist; orlistat, Orl) for 8 hours and assessed for PfAtg18 localization by live cell fluorescence microscopy. The panels for the indicated treatments represent GFP/PfAtg18 signal (Atg18), nucleic acid staining (Hoechst), bright field showing the boundaries of parasite and RBC (DIC), and the overlap of all three images (Merged). The %Representation column on the right indicates the %parasites showing that type of localization. The data is mean with SD (in the bracket) from three independent experiments. Compared to PfAtg18 localization in control parasites, treatment with E64 and pepstatin A produced a ring-like localization pattern and orlistat caused diffuse localization.

### Atg18 is crucial for parasite development

To determine the role of Atg18 during parasite development, we attempted to knock-out and knock-down PbAtg18. Multiple attempts to generate knock-out parasites were not successful, whereas knock-down parasites (PbAtg18-KD) were readily obtained by replacing the wildtype PbAtg18 coding sequence with PbAtg18/cDD_HA_ coding sequence (Figure 11A and B). cDD is a mutant *E. coli* DHFR, and cDD-fusion proteins undergo degradation in the absence of trimethoprim (TMP) but not in the presence of TMP, thereby producing a knock-down phenotype. The PbAtg18/cDD_HA_ fusion protein was localized to haemozoin-containing vesicles in PbAtg18-KD parasites (Figure 11C), which also showed slightly lower Atg18 level in the absence of TMP than the parasites grown in the presence of TMP (Figure 11D), indicating a partial knock-down effect. The effect of knock-down on development of PbAtg18-KD parasites was assessed in BALB/c and C57/BL/6J mice. Without TMP, PbAtg18-KD parasites showed drastically reduced growth and were eventually cleared, whereas wild type and PbAtg18-KD (with TMP) parasites grew similarly and the infected mice had to be euthanized or succumbed to high parasitemia (Figure 11E and F), indicating indispensability of PbAtg18 for parasite development.

**Figure 11.**
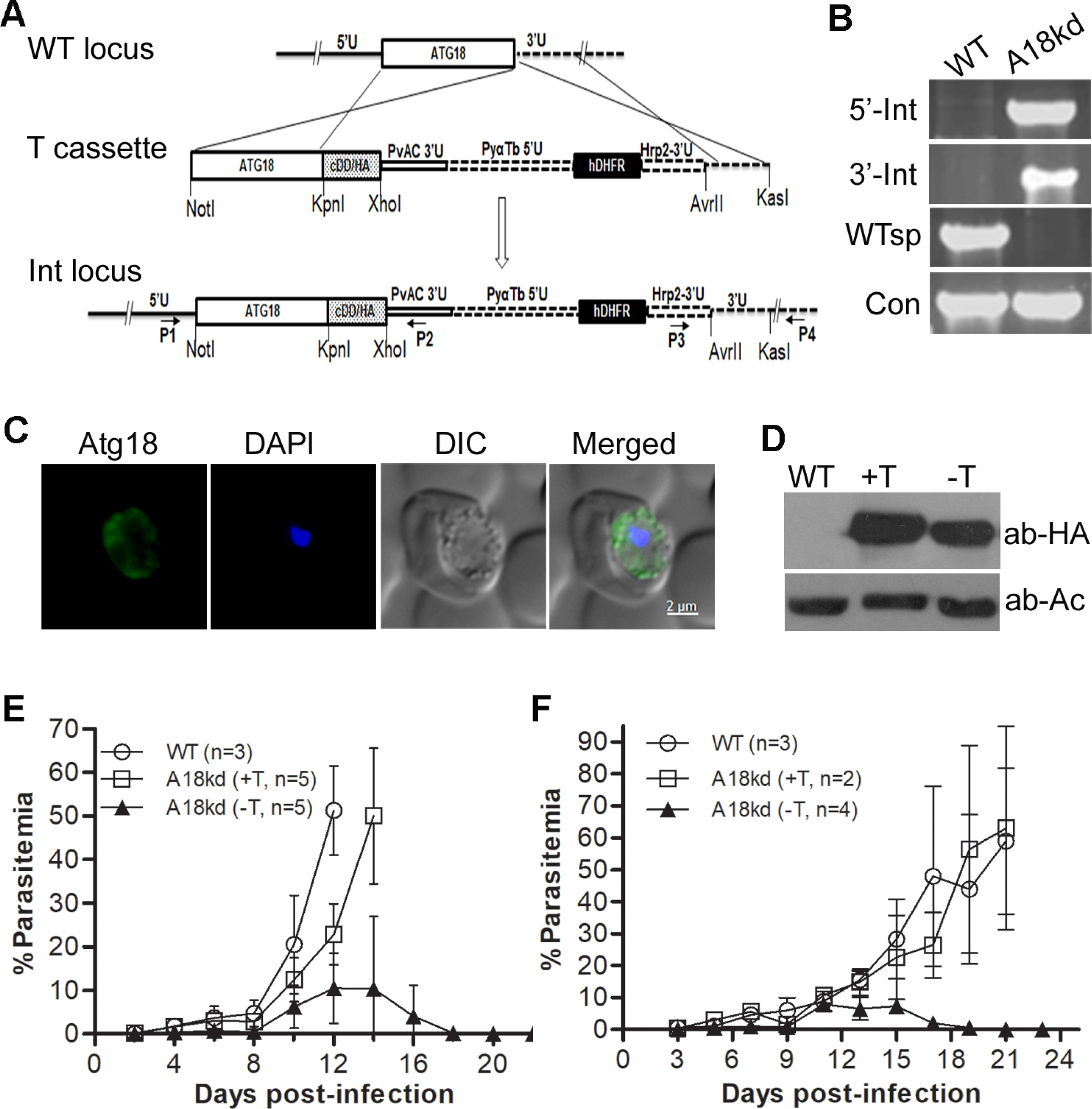
Atg18 is critical for erythrocytic stage parasite development. The endogenous PbAtg18 coding region was replaced with PbAtg18/cDD_HA_ coding sequence, and a clonal PbAtg18-KD parasite line was evaluated for the presence of knock-down locus by PCR, localization of PbAtg18-cDD/_HA_ protein, effect of knock-down on the expression of PbAtg18/cDD_HA_ protein level, and for comparision of growth rates with the wild type parasites. **A.** The schematic represents integration of the linear transfection construct (T cassette) into the endogenous PbAtg18 locus (WT locus), resulting into the generation of integration locus (Int locus). The coding regions for PbAtg18 (Atg18), cDD/HA, and hDHFR are represented by rectangular boxes. The flanking untranslated regions (5′U and 3′U), the location and orientation of primers (horizontal arrows), the restriction endonuclease sites, the regulatory regions in the linear transfection construct (PvAC3′U, PyαTb5′U and PfHrp2-3′U) are labelled. The wild type and integration loci can be distinguished by PCR using locus specific primers (5’ integration, 5’-Int (P1/P2); 3’ integration, 3’-Int (P3/P4); wild type locus, WTsp (P1/P4); MSP1 gene as a control, Con). **B.** The ethidium bromide stained agarose gel shows PCR products for the indicated regions from wild type (WT) and Atg18kd parasite gDNAs. **C.** The IFA images of PbAtg18-KD trophozoites show localization of Atg18/cDD_HA_ (Atg18), nucleic acid staining (DAPI), the bright field image showing boundaries of parasite and RBC (DIC), and the overlap of all three images (Merged). **D.** Western blot of WT and the PbAtg18-KD parasite lysates (grown in the presence (+T) or absence (-T) of trimethoprim) using antibodies to HA (ab-HA) and β-actin (ab-Ac). Naïve BALB/c **(E)** and C57BL/6J **(F)** mice were infected with equal number of wild type *P. berghei* ANKA (WT) or PbAtg18-KD (A18kd) parasites. The mice infected with PbAtg18-KD were kept under (A8kd, +T) or without (A18kd, -T) trimethoprim. The number of mice in each group is shown with “n”. The parasite growth is shown as %parasitemia over days post-infection.

## Discussion

*Plasmodium* Atg8 is present as punctate structures throughout the parasite, which also co-localize with the food vacuole and apicoplast [47, 48, 51]. Altered distribution of Atg8 puncta and mutation in PfAtg18 have been observed in chloroquine and artemisinin resistant *P. falciparum* strain*s*, respectively. We investigated *Plasmodium* Atg18 to understand the mechanism of its localization and association with drug resistance. Our study reveals that food vacuole localization is a conserved feature of *Plasmodium* Atg18 proteins, which is mediated by interaction with PI3P and MDR1 on the food vacuole membrane. Chloroquine and amodiaquine altered PfAtg18 localization in a significant number of cells. Food vacuole localization and interaction with MDR1 suggest that PfAtg18 could modulate the food vacuole-associated processes and function of MDR1.

PfAtg18 and its orthologs in *P. vivax*, *P. knowlesi* and *P. berghei* localized to the food vacuole, indicating that food vacuole localization is a conserved feature of *Plasmodium* Atg18 proteins. Interaction of PfAtg18 with the food vacuole-associated PI3P was critical for food vacuole localization, as PfAtg18 mutants of PI3P-binding sites (PIPm and ALCAm) had complete cytosolic localization. Based on the PfAtg18 homology model, “WLCL” motif forms part of the 23^rd^ β strand in blade 6, which contains the 2^nd^ PI3P-binding site and the residues contributing to the PI3P-binding site are highly conserved in PROPPINs [18, 23], suggesting that “WLCL” motif is a critical structural requirement for the formation of 2^nd^ PI3P-binding site in PfAtg18. The failure of ALCAm to bind PI3P and localize to the food vacuole could be due to disruption of the 2^nd^ PI3P-binding site in PfAtg18. Hence, the “WLCL” motif of PfAtg18 and the corresponding sequence in other PROPPINs represent a new sequence requirement for Atg18 functions. The PI3P-mediated localization of PfAtg18 in our study is consistent with previous reports of FRRG-mediated interaction of PROPPINs with PI3P, and the essentiality of this interaction for recruitment to membrane structures [18-23, 47, 48, 72].

The localization of PfAtg18 in our study is in agreement with recent reports [47, 48, 51], which showed association of PfAtg18 with the food vacuole and punctate structures all over the parasite. A slight difference in PfAtg18 localization between our and previous studies may be due to different procedures used for localization: live cell imaging in our study and IFA of fixed parasites in previous reports. IFA with anti-PfAtg18 antibodies did not work in our hands, possibly due to failure to recognize native PfAtg18 conformation, which is expected to be retained even in cells fixed with formaldehyde. Nonetheless, multiple lines of data support food vacuole localization of PfAtg18 in our study. 1) *P. falciparum* has been shown to have abundant PI3P in erythrocytic stages and localization studies using the PI3P probe GFP-2xFYVE indicated that PI3P is abundantly associated with the food vacuole membrane [44], which is consistent with PI3P-dependent localization of PfAtg18 to the food vacuole. 2) The localization was altered upon treatment of parasites with the PI3K inhibitor LY294002, which might be due to decreased production of PI3P, hence, less PI3P on the food vacuole membrane. 3) PIPm and ALCAm neither bound PI3P nor localized to the food vacuole. 4) In *P. berghei* that contains multiple food vacuoles, the episomally expressed GFP/PfAtg18 and endogenously expressed PbAtg18/GFP showed identical localization with food vacuoles, which ruled out any artefact of episomal expression and GFP fusion. 5) GFP fusions of ScAtg18 and WIPI have been extensively used in autophagy-related studies without any report of adverse effect on localization and functions of these proteins [18, 34, 72]. A previous study of the purified *P. falciparum* food vacuole proteome also contained PfAtg18 together with other food vacuole proteases and transporters, which further corroborates our result [87].

ScAtg18 has been shown to localize to endosomes, vacuole and autophagic membranes, and peri-vacuole membrane compartments in a PI3P-dependent manner [22, 29, 72, 88]. ScAtg18 also localizes on the vacuole membrane as a complex that regulates PI3P and PI3,5P2 levels on this organelle [40]. However, unlike *Plasmodium* Atg18 proteins investigated in this study, ScAtg18 showed complete cytoplasmic localization in *P. falciparum*, suggesting that interaction with PI3P is not the sole mediator for recruitment to the food vacuole. To identify the possible additional mediator of PfAtg18 localization, we carried out mass spectrometry of GFP/PfAtg18 and PbAtg18/GFP immunoprecipitates. MDR1, a food vacuole membrane protein [74], was reproducibly identified in both the immunoprecipitates and recombinant PfMDR1 interacted with recombinant PfAtg18. Recombinant ScAtg18 did not interact with recombinant MDR1, though both PfAtg18 and ScAtg18 interacted with PI3P, which could be a reason why ScAtg18 failed to localize to the food vacuole. ScAtg18 interacts with Atg2, and the Atg18-Atg2 complex is recruited to PAS through interaction of Atg18 with PI3P [34]. The apparent absence of Atg2 homolog in *P. falciparum* and differences in binding affinities of PfAtg18 and ScAtg18 for PI3P on the food vacuole membrane may also contribute to the failure of ScAtg18 to localize to the food vacuole. Hence, interaction with both PI3P and MDR1 is critical for PfAtg18 localization to the food vacuole.

The altered PfAtg18 localization upon chloroquine and amodiaquine treatment is of particular significance because quinolines, particularly chloroquine, has been proposed to block heme polymerization to hemozoin in the food vacuole and alter Atg8 distribution in chloroquine sensitive strains as compared to that in resistant strains of *P. falciparum* [12, 89, 90]. In mammalian cells, quinolines have been shown to inhibit autophagy by alkalinizing lysosomes, thereby inhibiting the activities of lysosomal hydrolases [91, 92]. However, the concentration required for alkalinization of the food vacuole is 1-2 orders of magnitude higher than the concentrations used in our study [90]. Moreover, treatment of parasites with NH_4_Cl, which inhibits vesicle fusion and lysosomal activity by alkalinizing these compartments [65], did not result in diffuse localization of PfAtg18. Also, treatment of parasites with mefloquine, halofantrine and quinine did not have any effect. Furthermore, E64 and chloroquine did not affect the food vacuole localization of PfCRTmCherry, a food vacuole membrane transport protein, indicating that food vacuole integrity was not compromised in our experiments. Hence, the diffuse localization of PfAtg18 in parasites treated with chloroquine and amodiaquine is unlikely due to alkalinization or disruption of the food vacuole, suggesting for an additional mechanism of the action of chloroquine and amodiaquine at the level of Atg18 recruitment to the food vacuole, which may contribute to overall antimalarial effect of these compounds. ScAtg18 and PfAtg18 have also been shown to be involved in vacuole and food vacuole dynamics, respectively [41, 42, 48], suggesting that altered food vacuole localization of PfAtg18 could affect multiple food vacuole-associated processes. Further investigation is necessary whether chloroquine and amodiaquine directly target PfAtg18 or additional proteins with a role in PfAtg18 localization.

The *Plasmodium* Atg18 gene is essential for parasite development [93, 94], and knock-down of PfAtg18 has been shown to cause loss of apicoplast [47]. We generated PbAtg18-KD parasites, which did not show any noticeable change in the localization of PbAtg18, most likely because there was only a partial reduction in the PbAtg18 protein level. A more efficient knock-down approach will be required to define the role of *Plasmodium* Atg18 in autophagy and the functional importance of its association with the food vacuole.

In addition to the Cvt and autophagy pathways, Atg18 proteins of *S. cerevisiae* and *P. pastoris* are also required for pexophagy. The proteins are concentrated at one or more spots that resemble protuberances of the vacuole membrane [31]. Elegant microscopic images indicated colocalization of the protuberance with peroxisome cluster during micropexophagy [31, 95]. PfAtg18 also showed an intense spot on or near the food vacuole membrane, which tempted us to speculate if PfAtg18 functions in engulfment of cellular contents directly by the food vacuole membrane. Although peroxisomes appear to be absent in *Plasmodium* [96], direct engulfment of cellular contents by the *Plasmodium* food vacuole membrane may occur. One such process could be the fusion of haemoglobin-containing vesicles with the food vacuole [97, 98]. Similar PfAtg18-enriched dot-like structure in the food vacuole periphery was also observed in a recent study, which the authors explained as a result of the food vacuole fission [48]. ScAtg18 has also been shown to be present as an intense spot in a peri-vacuole membrane compartment during starvation wherein it colocalizes with a number of autophagy proteins (Atg1, Vps34, Atg8, Atg9), and this compartment has been proposed to be a site for vesicle formation or membrane source [43]. The intense PfAtg18 spot in the peri-food vacuole region may be a platform where autophagy and some other pathways overlap. As many proteins required for autophagosome biogenesis (Atg2, Atg10 and Atg16) remain to be identified in *Plasmodium*, further studies are required to gain insights into the nature and function of this peri-food vacuole compartment.

Thus, our data demonstrate that food vacuole localization is conserved among *Plasmodium* Atg18 proteins and it is mediated by interaction with PI3P and MDR1. Since single nucleotide polymorphisms and increased copy number of MDR1 are associated with resistance to multiple antimalarials [49, 75–78], it is possible that PfAtg18 modulates MDR1 activity, including the associated drug resistance. Additional study is needed to investigate the physiological significance of MDR1-PfAtg18 interaction.

## Data availability

The mass spectrometry proteomics data have been deposited to the ProteomeXchange Consortium via the PRIDE partner repository with the dataset identifier PXD020847 and 10.6019/PXD020847.

## Acknowledgements

We thank the staff of CCMB’s Laboratory Animal Facility, Advanced Microscopy and Imaging Facility, and Proteomics Facility for assistance with experiments related to mice, imaging, and mass spectrometry, respectively. We thank Dr. Radhika Khandelwal for assistance with the CD experiments.

## Funding information

This study was supported by grants from the Department of Biotechnology, India (BT/PR11497/MED/29/854/2014) and Council of Scientific and Industrial Research, India (MLP0148) to PS. The salary of DD was supported by a grant from the Department of Biotechnology, India (BT/PR11497/MED/29/854/2014). RS is a recipient of fellowships from the Department of Biotechnology, India.

## Conflict of interest

The authors do not have any conflict of interest with the contents of this article.

---

**Table Sl.**
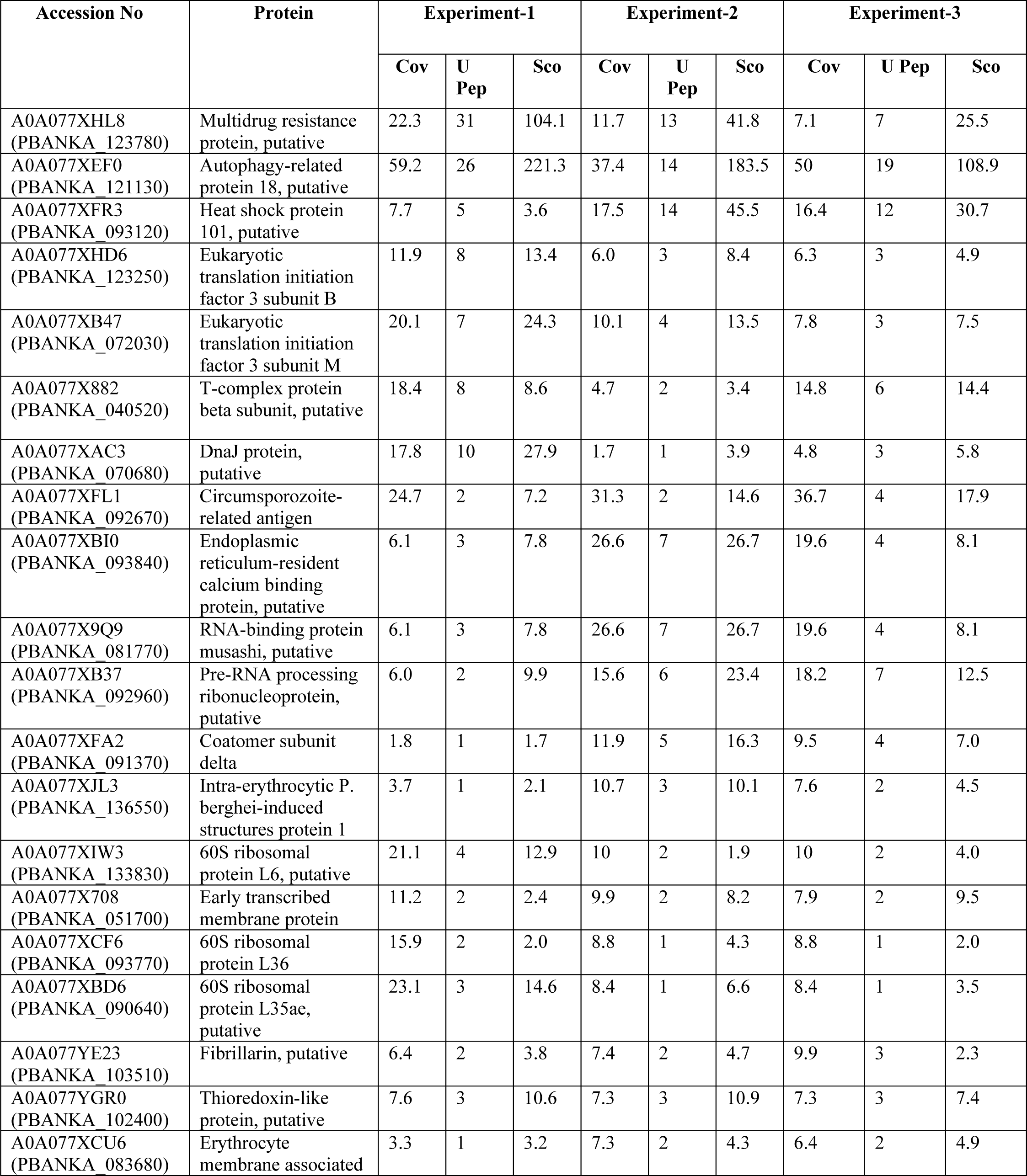
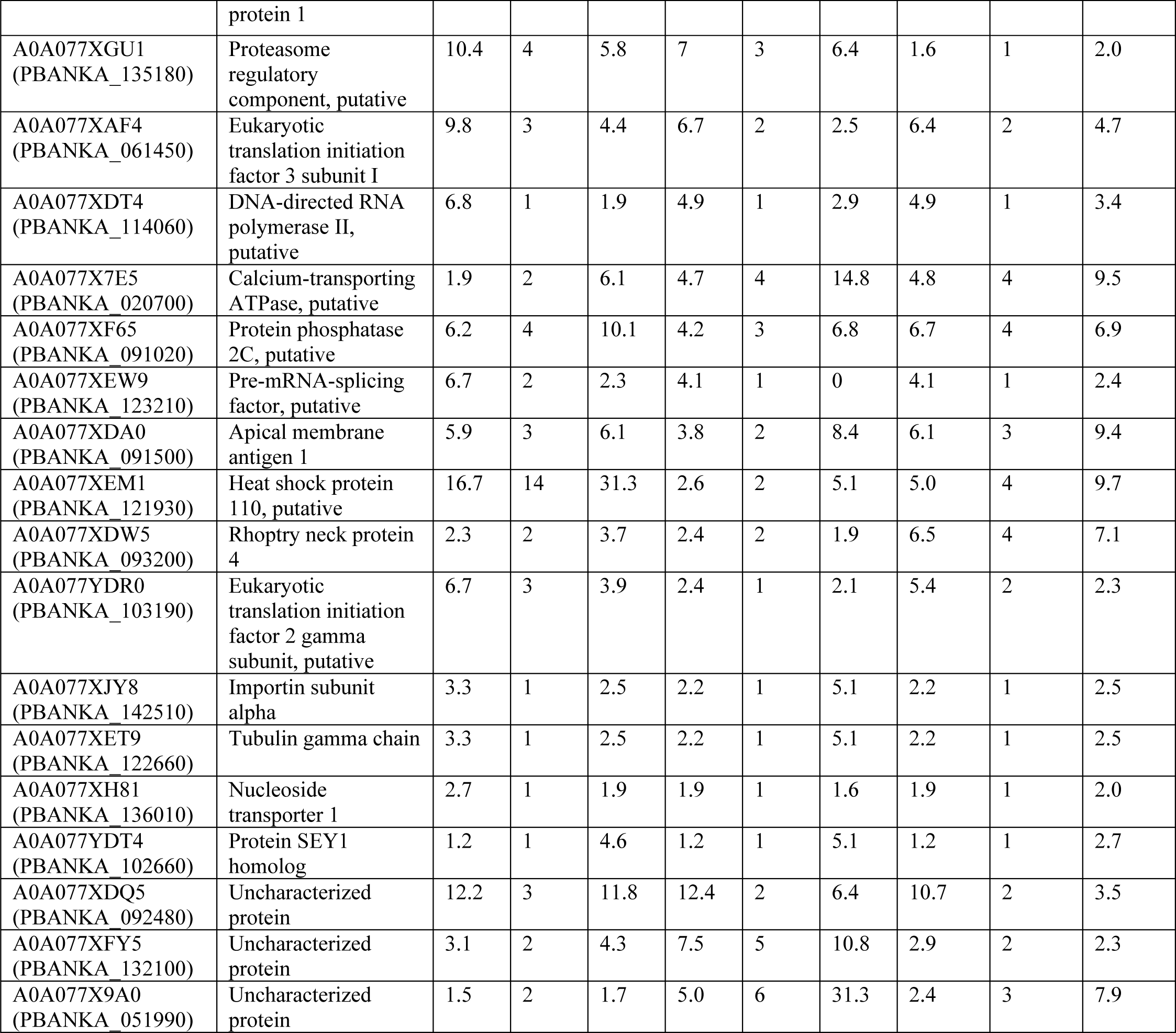
Proteins identified in the immunoprecipitate of PbAtg18/GFP-expressing *P. berghei* parasites. The proteins listed are common in at least 3 independent experiments, and contain at least 1 unique peptide. Shown are coverage (Cov), unique peptides (U Pep) and score (Sco) for each protein.

**Table S2.**
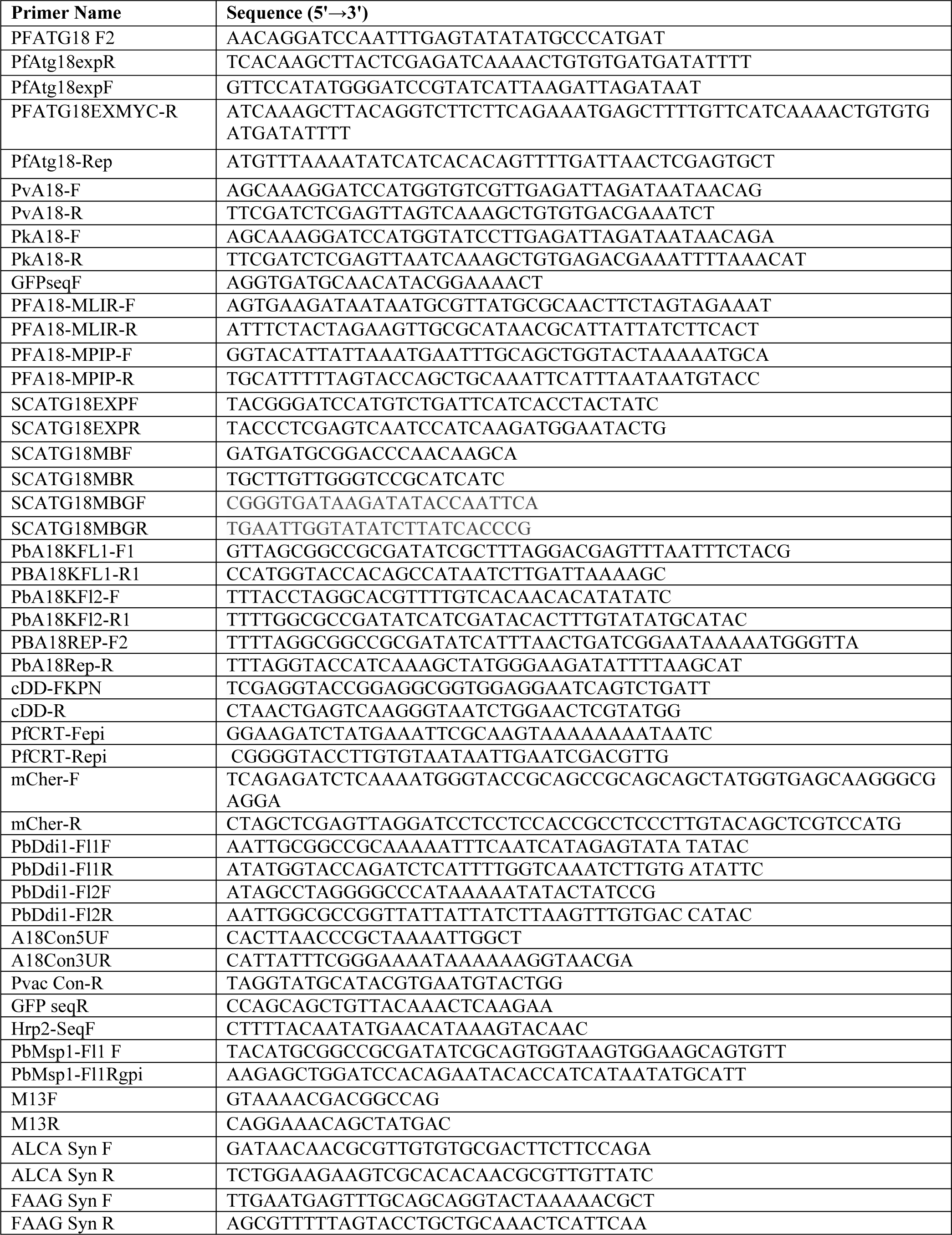
List of primers used in the study.

**Table S3.** IC_50_ concentrations of inhibitors and antimalarials. The table shows IC_50_ values of different inhibitors and antimalarials, which were determined using 48-hour parasite survival assay using *P. falciparum* as has been described previously (1). The IC_50_ for compounds represented with “*” are taken from the referenced reports (1-4).

| Inhibitors | IC<sub>50</sub> for 3D7 | Concentrations used in the study |
| --- | --- | --- |
| E64 <sup>*1</sup> | 7.6 μM | 22 μM |
| Pepstatin <sup>*(1)</sup> | 74.3 μM | 300 μM |
| Epoxomicin <sup>*1</sup> | 7.7 nM | 30.8 nM |
| Orlistat | 13.1 μM | 52.4 μM |
| Pristemerin | 0.12 μM | 0.48 μM |
| Chloroquine | 27 nM | 120 nM |
| Artemisinin | 59.1 nM | 236.4 nM |
| Amodiaquine <sup>*2</sup> | 20.4 nM | 81.6 nM |
| Halofantrine <sup>*2</sup> | 5.4 nM | 21.6 nM |
| Quinine <sup>*2</sup> | 133 nM | 532 nM |
| Mefloquine <sup>*2</sup> | 31.2 nM | 124.8 nM |
| LY294002 | 31.3 μM | 95 μM |
| Bafilomycin <sup>*4</sup> | 21-26 nM | 75 nM and 100 nM |
| Brefeldin A <sup>*3</sup> | Not determined | 20 μM |
| NH<sub>4</sub>Cl <sup>*2</sup> | 3.4 mM | 13.6 mM |

**Figure S1.**
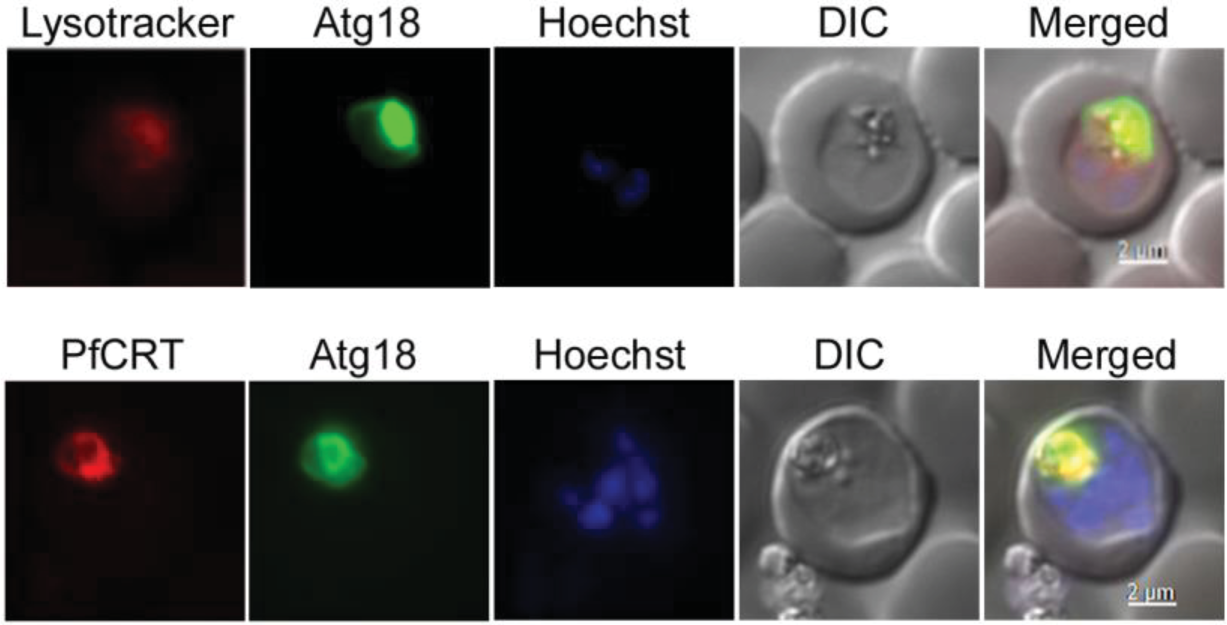
Colocalization of PfAtg18 with food vacuole markers. GFP/PfAtg18-expressing *P. falciparum* trophozoites cultured in the presence of lysotracker, and *P. falciparum* trophozoites co-expressing GFP/PfAtg18 and PfCRT/mCherry were assessed by live cell fluorescence microscopy. Panels from the left represent signal for lysotracker or PfCRT, GFP/PfAtg18 (Atg18), nucleic acid staining (Hoechst), bright field showing parasite and RBC boundaries (DIC), and the merge of all images (Merged). Both lysotracker and PfCRT overlap with GFP/PfAtg18, and also colocalize with the in-built food vacuole marker haemozoin.

**Figure S2.**
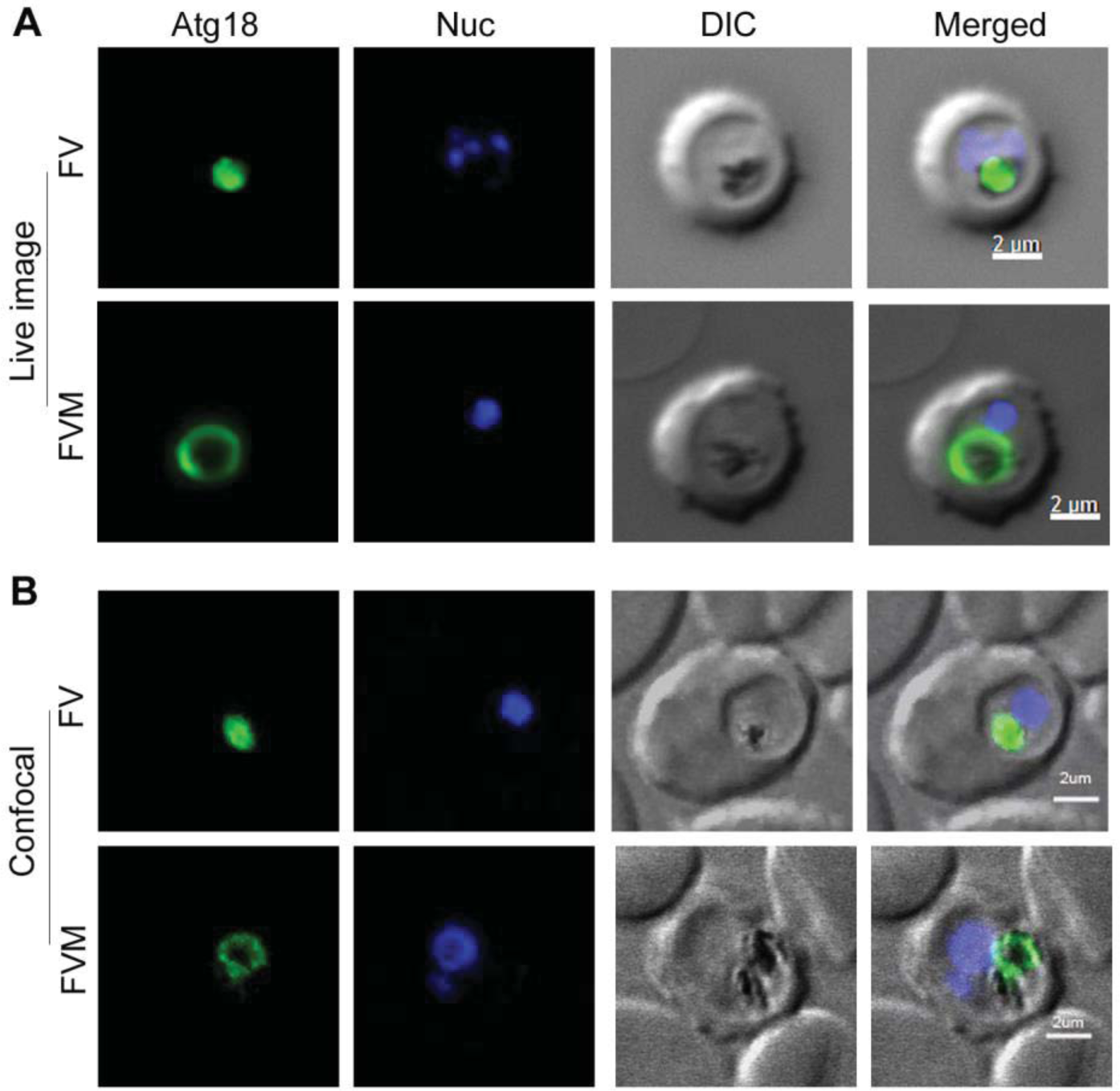
Localization patterns of PfAtg18. Live or fixed trophozoites of GFP/PfAtg18-expressing parasites were evaluated for localization of PfAtg18 by live cell fluorescence microscopy (**A**) and confocal microscopy (**B**), respectively. The images are representative for PfAtg18 within the food vacuole (FV) and around the food vacuole membrane (FVM). Panels from the left represent signals for GFP/PfAtg18 (Atg18), nucleic acid staining (Nuc), bright field with parasite and RBC boundaries (DIC), and the merge of all images (Merged).

**Figure S3.**
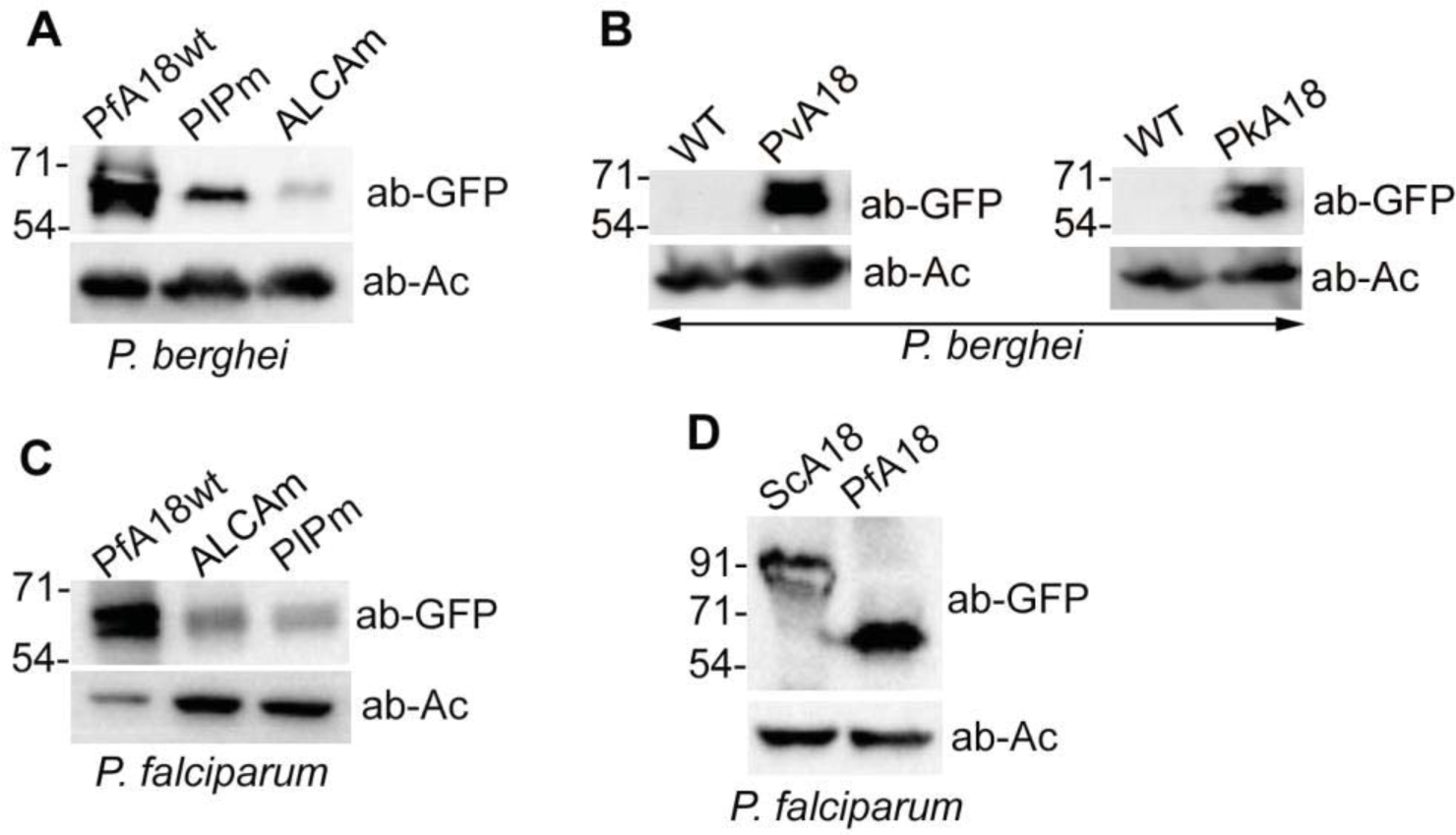
Western blots of parasites expressing different Atg18 proteins. Trophozoite stages of wild type (WT) and recombinant parasites were assessed for expression of different Atg18 proteins using anti-GFP antibodies (ab-GFP). β-Actin expression was detected using β-actin antibodies (ab-Ac) and used as a loading control. **A** and **C** show expression of wild type GFP/PfAtg18 (PfA18wt) and its mutants (PIPm and ALCAm) in *P. berghei* and *P. falciparum* D10 parasites. **B.** The western blot shows expression of GFP/PvAtg18 (PvA18) and GFP/PkAtg18 (PkA18) in *P. berghei*. **D.** The western blot shows expression of GFP/ScAtg18 (ScA18) and GFP/PfAtg18 (PfA18) in *P. falciparum* 3D7 parasites. The blots correspond to regions containing the expected proteins and the numbers on the left side of each blot indicate the sizes of protein molecular mass markers in kDa. The antibodies only or predominantly detected the expected proteins, which were absent in lanes of wild type parasite samples.

**Figure S4.**
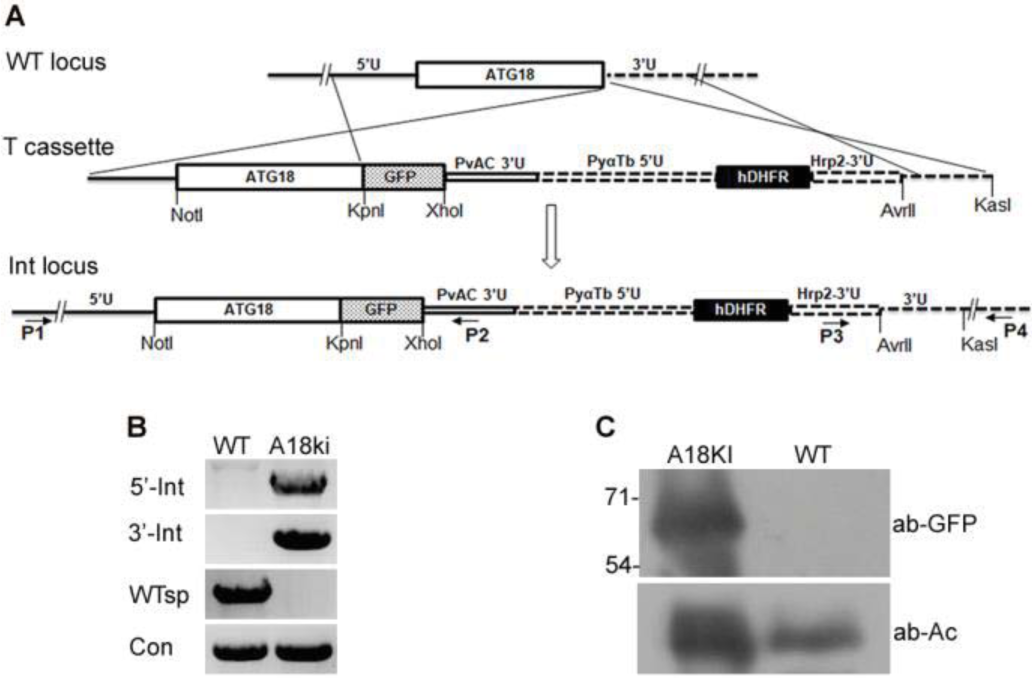
Generation of PbAtg18/GFP knock-in parasites. The endogenous PbAtg18 coding sequence was replaced with PbAtg18/GFP coding sequence. Recombinant parasites were evaluated for replacement of the wild type locus by PCR and expression of PbAtg18/GFP protein by western blotting. **A**. The schematic represents replacement of the endogenous PbAtg18 locus (WT locus) by a linear transfection construct (T cassette), generating the integration locus (Int locus). The coding regions for PbAtg18 (Atg18), GFP and hDHFR are represented by rectangular boxes. The flanking untranslated regions (5′U and 3′U), the location and orientation of primers (horizontal arrows), the restriction endonuclease sites, the regulatory regions in the linear transfection construct (PvAC3′U, PyαTb5′U and PfHrp2-3′U) are labelled. The wild type and integration loci can be distinguished by PCR using locus specific primers (5’ integration, 5’-Int (P1/P2); 3’ integration, 3’-Int (P3/P4); wild type locus, WTsp (P1/P4)). Primers for MSP1 gene were used as a control (Con). **B.** The ethidium bromide stained agarose gel shows PCR products for indicated regions from the wild type (WT) and PbAtg18/GFP-KI (A18ki) parasite gDNAs. **C.** Western blot of WT and A18ki parasite lysates using antibodies to GFP (ab-GFP) and β-actin (ab-Ac) as a loading control. The absence of wild type PbAtg18 gene and the presence of Atg18/GFP protein in PbAtg18/GFP-KI parasite samples confirmed the knock-in.

**Figure S5.**
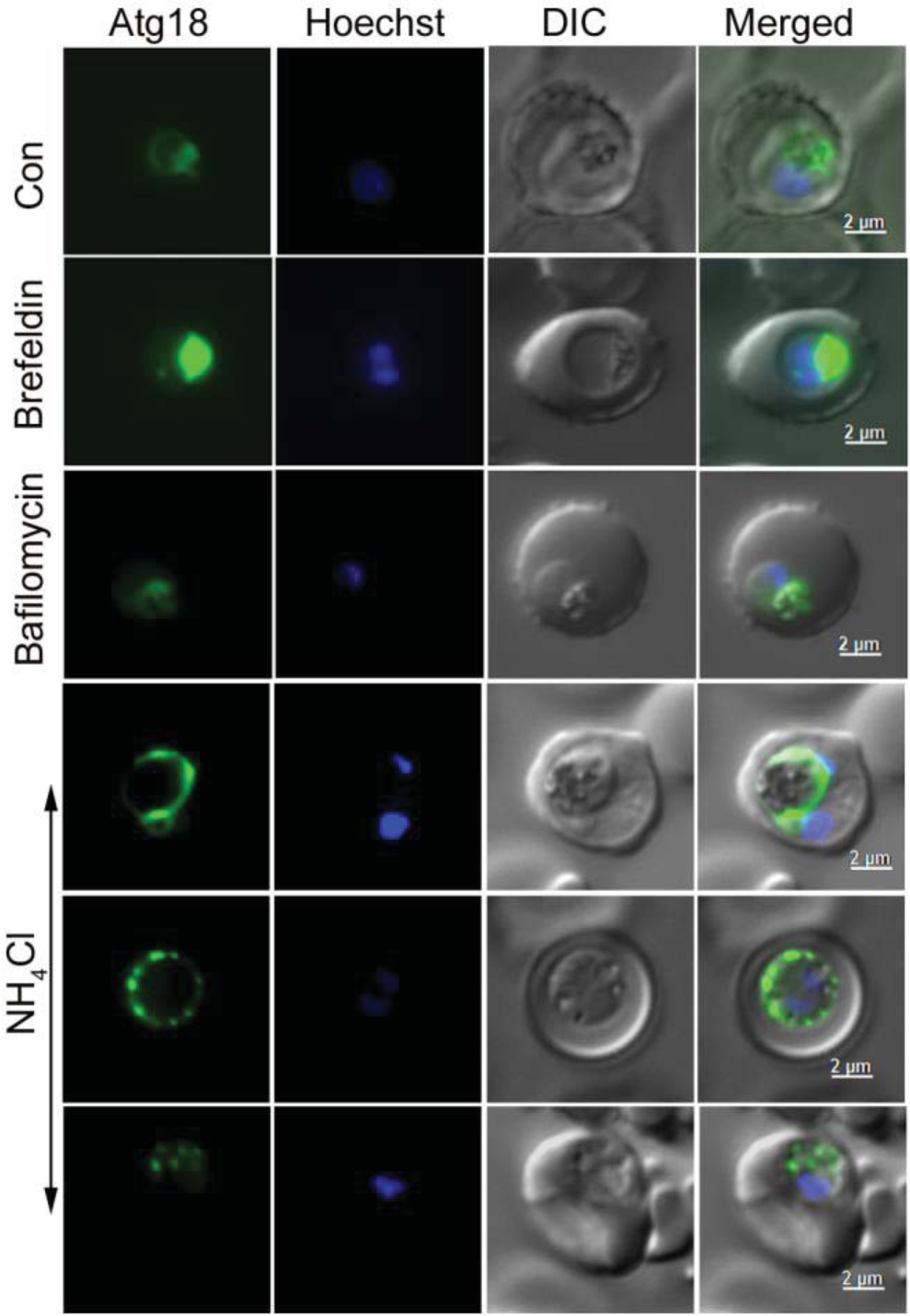
PfAtg18 localization upon treatment with endosomal transport inhibitors. GFP/PfAtg18-expressing *P. falciparum* parasites at late ring stage were cultured in the absence (Con) or presence of indicated endosomal transport inhibitors for 8 hours, and processed for live cell fluorescence microscopy. Shown are the representative images for GFP/PfAtg18 localization (Atg18), nucleic acid staining (Hoechst), bright field with parasite and RBC boundaries (DIC), and the merge of all images (Merged). As opposed to the food vacuole-associated signal of PfAtg18 in control parasites, NH_4_Cl-treated parasites show PfAtg18 around the enlarged food vacuole or as multiple foci near the parasite periphery or throughout the parasite.

**Figure S6.**
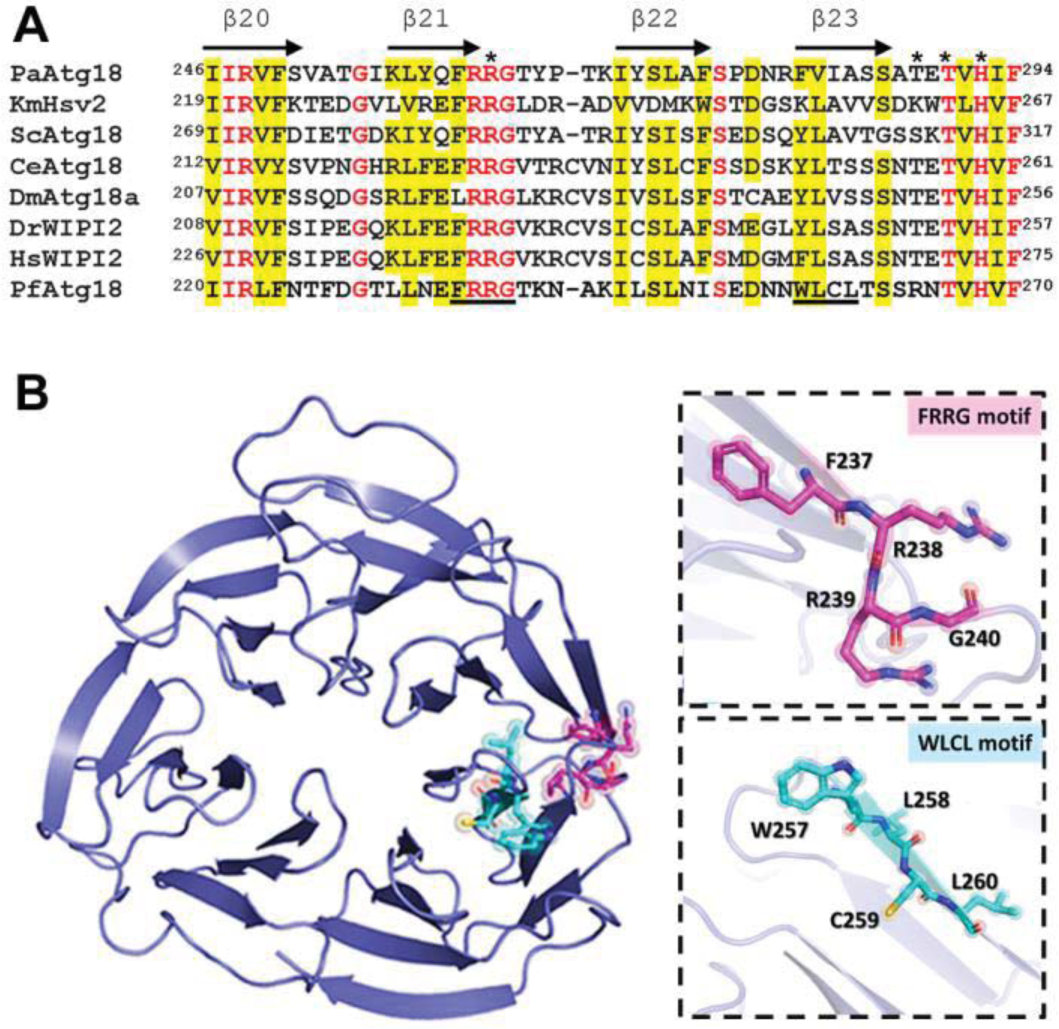
PfAtg18 is a PROPPIN family protein. **A.** Structure-based sequence alignment of *P. falciparum* Atg18 (PfAtg18) with the homologs from *Pichia angusta* (PaAtg18), *Kluyveromyces marxianus* (KmHsv2), *Saccharomyces cerevisiae* (ScAtg18), *Caenorhabditis elegans* (CeAtg18), *Drosophila melanogaster* (DmAtg18a), *Danio rerio* (DrAtg18) and *Homo sapiens* (HsWIPI2). The alignment corresponds to the region containing “FRRG” and “WLCL” motifs of PfAtg18. The conserved amino acids are in red font, physicochemically similar amino acids in 6 of the 8 sequences are shaded yellow, the motifs are underlined, and amino acids corresponding to the ScAtg18 2^nd^ PI3P-binding site are marked with asterisk. Numbers in the beginning and the end of each sequence indicate the position of corresponding residues in the sequence. The β strands of PaAtg18 are indicated with arrows on top of the sequence. **B.** The homology model of PfAtg18 on the left side shows its β-propeller structure, with the “FRRG” (magenta) and “WLCL” (cyan) motifs zoomed in on the right.

**Figure S7.**
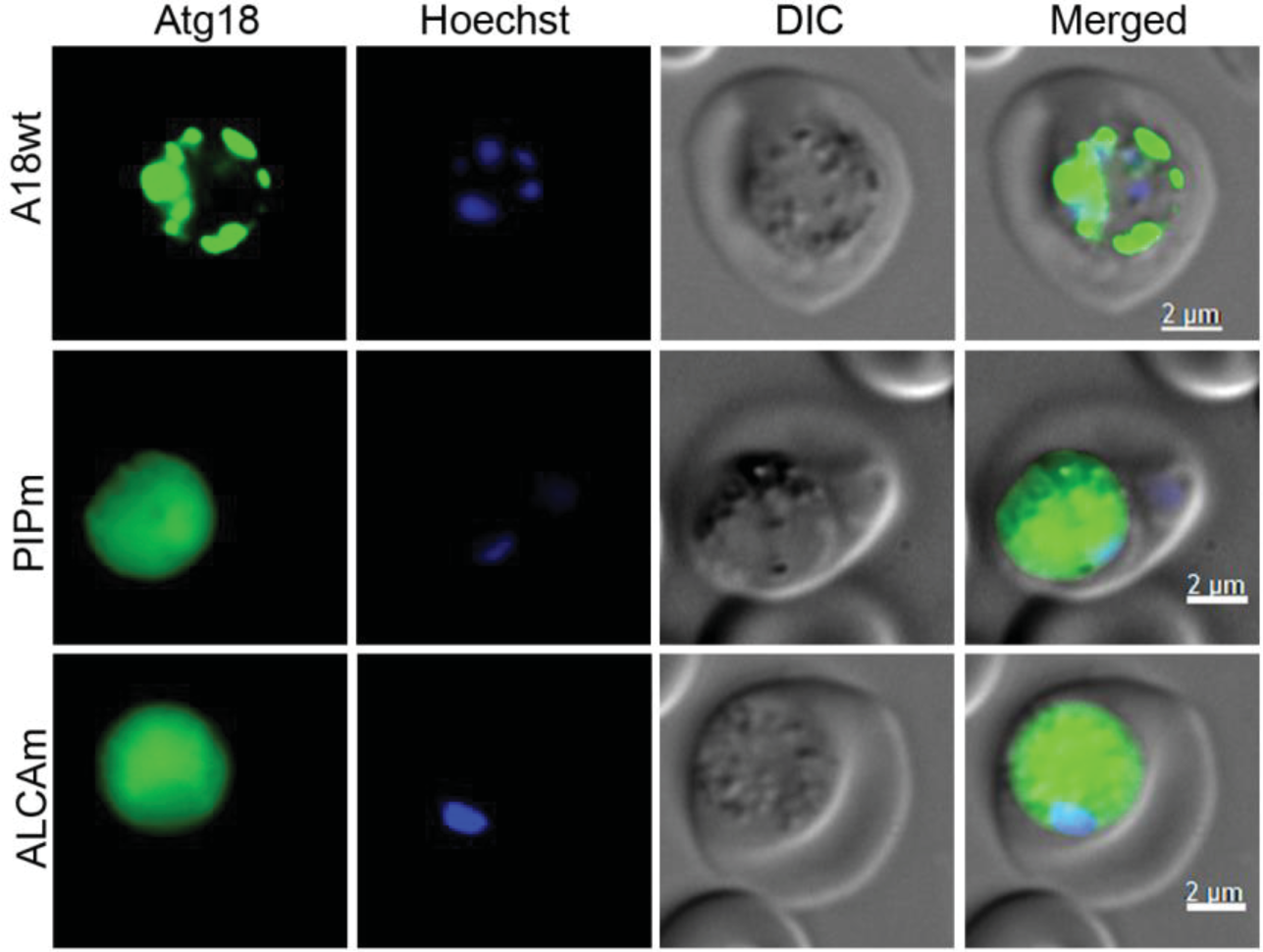
Localization of PfAtg18 mutants in *P. berghei*. Trophozoites of *P. berghei* expressing wild type GFP/PfAtg18 (A18wt), GFP/PIPm (PIPm) or GFP/ALCAm (ALCAm) were observed by live cell fluorescence microscopy. The panels from the left indicate signal for wild type PfAtg18 or its mutants (Atg18), nucleic acid staining (Hoechst), the boundaries of parasite and RBC (DIC), and the overlap of all three images (Merged). Both PIPm and ALCAm show diffuse localization all over the parasite, which is in contrast with the predominant food vacuole-associated localization of wild type PfAtg18.

**Figure S8.**
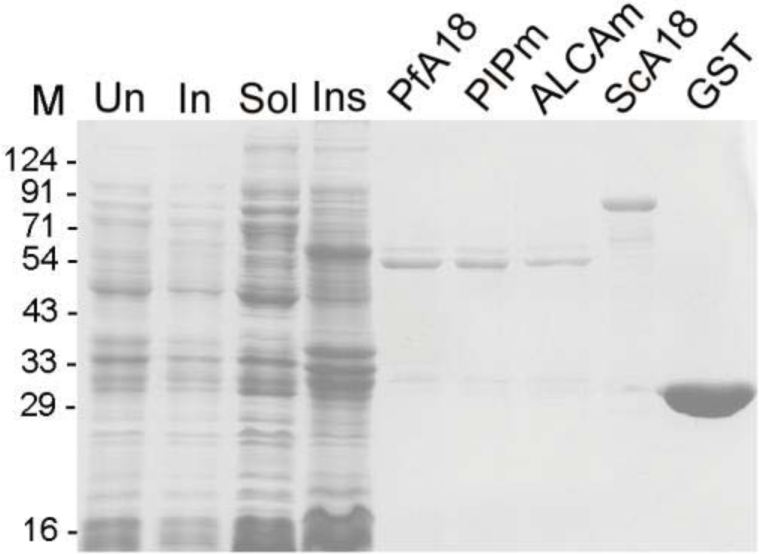
Production of GST-fusion proteins. The GST/PfAtg18syn (PfA18), GST/PIPm (PIPm), GST/ALCAm (ALCAm), GST/ScAtg18 (ScA18) and GST were expressed in BL21(DE3) *E. coli* cells, and purified from IPTG-induced cells. The coomassie-stained SDS-PAGE gel shows expression and purification of GST/PfAtg18syn. The lanes contain uninduced (Un) and induced (In) total cell lysates of GST/PfAtg18-expressing cells, soluble (Sol) and insoluble (Ins) extracts of the induced GST/PfAtg18-expressing cells, and the indicated purified proteins.

**Figure S9.**
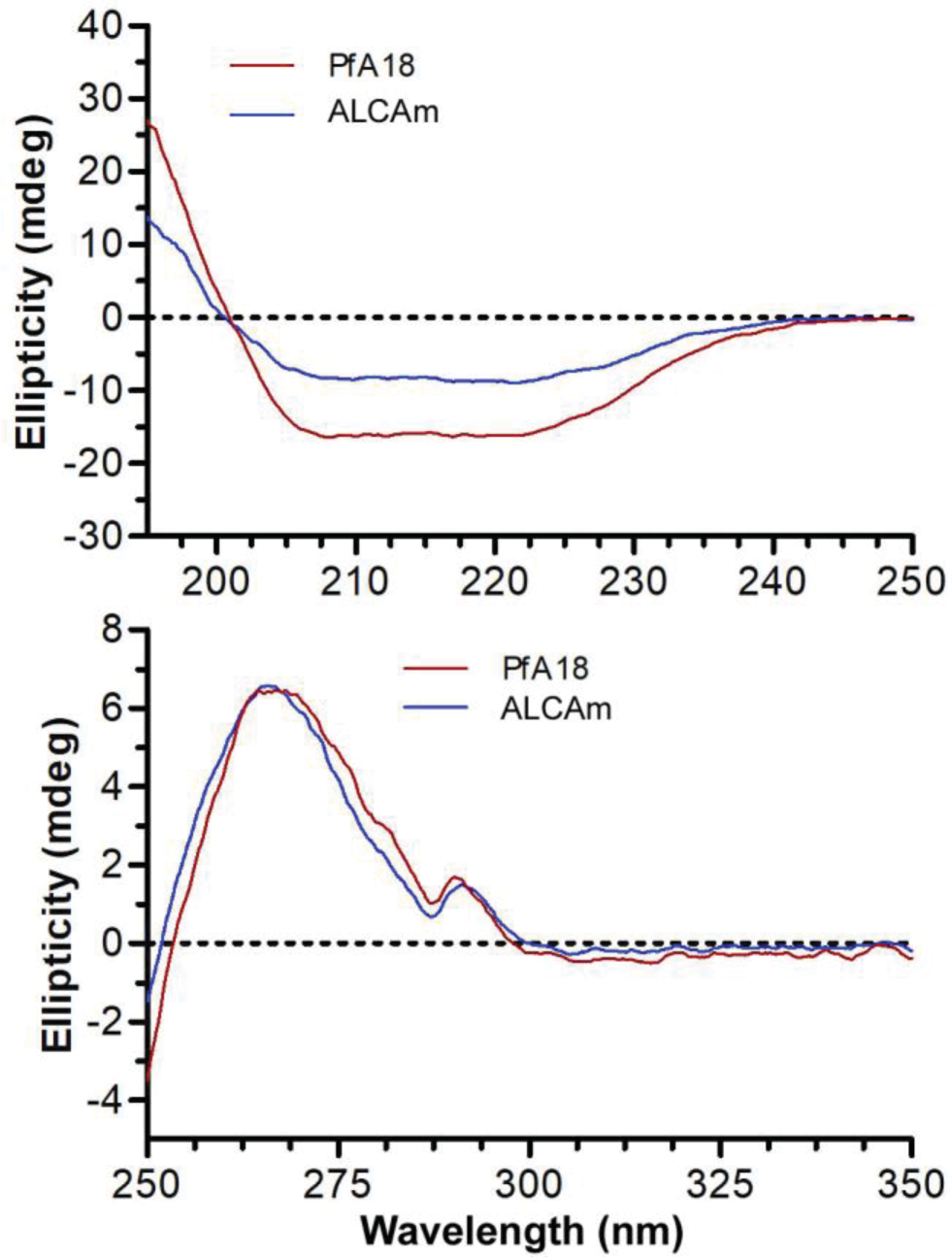
CD spectra of recombinant wild type and mutant PfAtg18. The far-UV and near-UV CD spectra of wild type GST/PfAtg18 (PfA18) and GST/ALCAm (ALCAm) proteins were recorded and molar ellipticity was calculated. The plots show mean residue molar ellipticity against the wavelength for the indicated proteins, and the CD spectra are an average of five accumulations.

**Figure S10.**
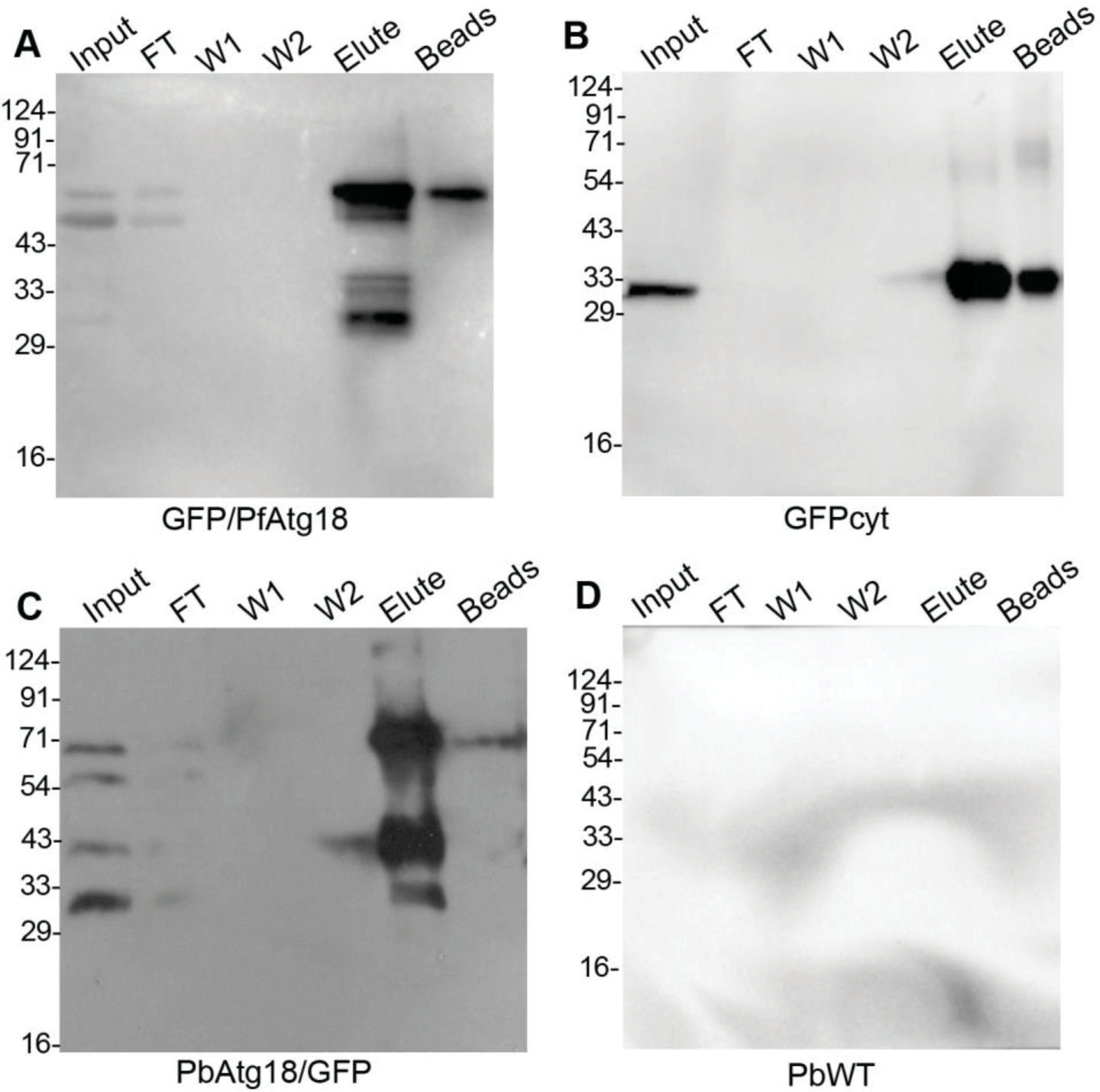
Immunoprecipitation of *Plasmodium* Atg18. The extracts of GFP/PfAtg18-expressing *P. falciparum* (GFP/PfAtg18, A), cytosolic GFP-expressing *P. falciparum* (GFPcyt, B), PbAtg18/GFP knock-in *P. berghei* (PbAtg18/GFP, C) and wild type *P. berghei* ANKA (PbWT, D) were incubated with GFP-Trap magnetic agarose resin. The resin was washed and bound proteins were eluted. The eluates (Elute) along with the extracts (Input), flow through (FT), washes (W1 and W2), and beads after elution were analysed by western blotting using rabbit anti-GFP antibodies. The western blots show the presence or absence of the target protein in indicated parasite samples. The protein size markers are in kDa (M).

**Figure S11.**
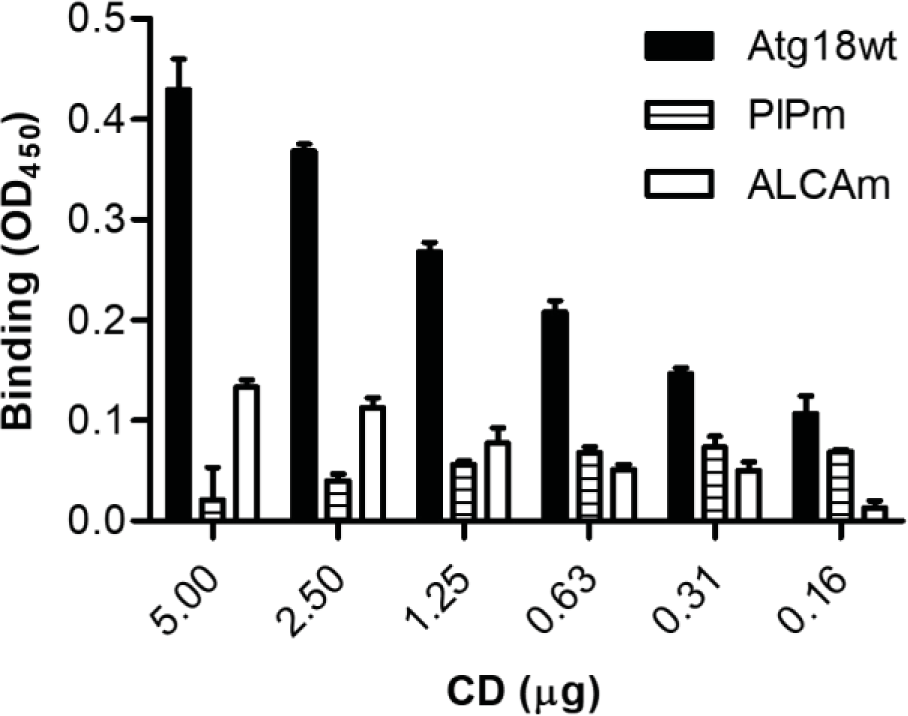
Interaction of wild type and mutant PfAtg18 proteins with MDR1. Equimolar concentrations of recombinant GST/PfAtg18syn (Atg18wt), GST/PIPm (PIPm), GST/ALCAm (ALCAm) or GST proteins were coated to the wells of an ELISA plate. The indicated concentrations of myc-tagged GST/MDRCD (CD) were incubated with the coated recombinant proteins, and the bound myc-tagged GST/MDRCD was detected using mouse anti-myc antibodies, followed by measuring absorbance at 450 nm as described in the experimental procedures. The plot shows the mean of background-adjusted absorbance as binding at different GST/MDRCD concentrations.

**Figure S12.**
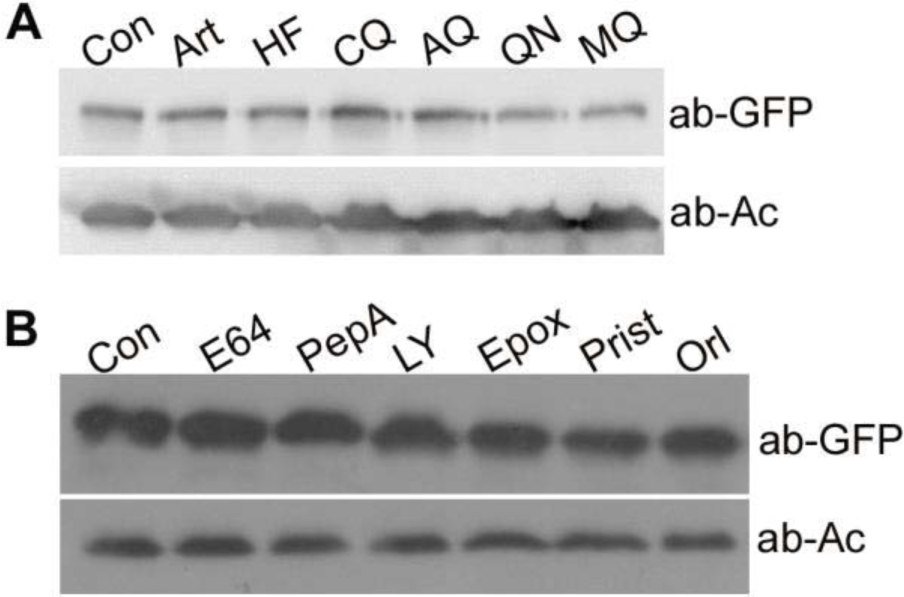
Effect of antimalarials and inhibitors on PfAtg18 expression. Highly synchronized cultures of GFP/PfAtg18-expressing *P. falciparum* parasites at early trophozoite stage were cultured in the presence of DMSO (Con) or (A) antimalarials (artemisinin, Art; halofantrine, HF; chloroquine, CQ; amodiaquine, AQ; quinine, QN; mefloquine, MQ) or (B) inhibitors (E64; pepstatin A, PepA; LY294002, LY; epoxomicin, Epox; pristimerin, Prist; orlistat, Orl) for 8 hours, and processed for evaluation of PfAtg18 level by western blotting using anti-GFP antibodies (ab-GFP). Antibodies to β-actin (ab-Ac) were used to detect β-actin in the samples as a loading control. PfAtg18 levels were similar in control and the parasites treated with antimalarial or inhibitors.

**Figure S13.**
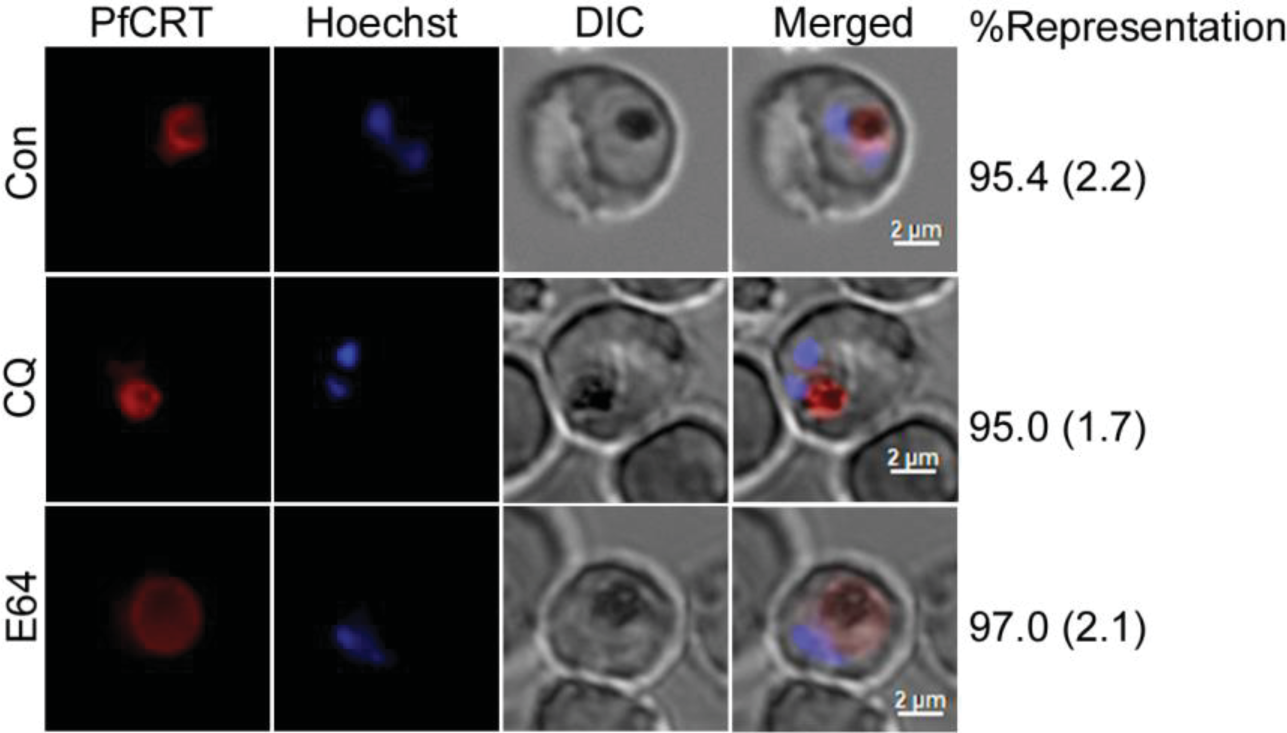
PfCRT localization upon treatment with the inhibitors of food vacuole-associated processes. Early trophozoite stage GFP/PfCRTmcherry-expressing parasites were treated with DMSO (Con), chloroquine (CQ) or E64 for 8 hours and assessed for PfCRT localization by live cell fluorescence microscopy. The panels for the indicated treatments represent GFP/PfCRTmcherry signal (PfCRT), nucleic acid staining (Hoechst), bright field showing the boundaries of parasite and RBC (DIC), and the overlap of all three images (Merged). The %Representation column on the right indicates the %parasites showing that type of localization, which is comparable in control and inhibitor treated parasites. The expansion of PfCRT localization in E64-treated parasite is due to enlargement of the food vacuole. The data is mean with SD (in the bracket) from three independent experiments.

## Notes

### Competing Interest Statement

The authors have declared no competing interest.

### Summary of Updates

Some of the data has been excluded from the manuscript, and text has minor edits.

